# HHIP protein interactions in lung cells provide insight into COPD pathogenesis

**DOI:** 10.1101/2024.04.01.586839

**Authors:** Dávid Deritei, Hiroyuki Inuzuka, Peter J. Castaldi, Jeong Hyun Yun, Zhonghui Xu, Wardatul Jannat Anamika, John M. Asara, Feng Guo, Xiaobo Zhou, Kimberly Glass, Wenyi Wei, Edwin K. Silverman

## Abstract

Chronic obstructive pulmonary disease (COPD) is the third leading cause of death worldwide. The primary causes of COPD are environmental, including cigarette smoking; however, genetic susceptibility also contributes to COPD risk. Genome-Wide Association Studies (GWASes) have revealed more than 80 genetic loci associated with COPD, leading to the identification of multiple COPD GWAS genes. However, the biological relationships between the identified COPD susceptibility genes are largely unknown. Genes associated with a complex disease are often in close network proximity, *i.e.* their protein products often interact directly with each other and/or similar proteins. In this study, we use affinity purification mass spectrometry (AP-MS) to identify protein interactions with *HHIP*, a well-established COPD GWAS gene which is part of the sonic hedgehog pathway, in two disease-relevant lung cell lines (IMR90 and 16HBE). To better understand the network neighborhood of *HHIP*, its proximity to the protein products of other COPD GWAS genes, and its functional role in COPD pathogenesis, we create HUBRIS, a protein-protein interaction network compiled from 8 publicly available databases. We identified both common and cell type-specific protein-protein interactors of HHIP. We find that our newly identified interactions shorten the network distance between HHIP and the protein products of several COPD GWAS genes, including *DSP, MFAP2, TET2*, and *FBLN5*. These new shorter paths include proteins that are encoded by genes involved in extracellular matrix and tissue organization. We found and validated interactions to proteins that provide new insights into COPD pathobiology, including CAVIN1 (IMR90) and TP53 (16HBE). The newly discovered HHIP interactions with CAVIN1 and TP53 implicate HHIP in response to oxidative stress.

## Introduction

Chronic obstructive pulmonary disease (COPD), a complex disease primarily caused by a combination of cigarette smoke exposure and genetic predisposition, is a leading cause of death in the United States and worldwide(1). Although clinically defined by airflow obstruction measured by spirometry, COPD is a heterogeneous syndrome that manifests with variable amounts of airway disease and/or destruction of the lung alveolar structures (emphysema). COPD is also associated with chronic inflammation(2, 3), remodeling of the extracellular matrix (ECM)(4), mitochondrial damage(5, 6), and cellular senescence(7, 8). However, the relative importance and timing of these potential pathomechanisms are not yet well understood. Characterizing the biological impact of the genetic determinants that predispose a person to COPD may provide key insights into the underlying mechanisms of COPD pathogenesis.

Genome-wide association studies (GWASes) have identified more than 80 genomic regions associated with COPD(9–11). In addition, based on a combination of genetic and functional evidence, the likely key genes have been identified for multiple COPD GWAS loci, including *HHIP*(*12*), *FAM13A*(*13*), *IREB2*(*6*), *DSP(14)*, *AGER(15)*, *MFAP2(16)*, *FBLN5(17)*, *NPNT(17)*, *FBXO38(18)*, *SFTPD(19)*, *TET2(20)*, *TGFB2(21)*, *MMP12(22)*, and *MMP1(23)*. Many of these genes have known functions outside of their role in COPD pathogenesis. For example, *HHIP*, the gene encoding the hedgehog interacting protein (previously referred to as HIP), was discovered as part of the Hedgehog pathway, where it acts as an endogenous inhibitor of the hedgehog proteins SHH, IHH, and DHH(24). *HHIP* was the first gene linked to COPD through GWAS, and genetic variants near *HHIP* have been consistently found to be associated with COPD. *HHIP*’s effects on COPD susceptibility have been confirmed by murine and human cell line experiments(25). For instance, *Hhip* haploinsufficient (+/-) mice exposed to chronic cigarette smoke are more likely to develop emphysema than wild type (+/+) mice(26, 27), while *Hhip* knockout (-/-) leads to faulty lung development and subsequent death(28). Bioinformatic evidence suggests that there may be functional interactions between *HHIP* and other COPD GWAS genes, but the nature of these connections is unknown(29).

While studying biological systems network analysis is a powerful tool that can reveal direct interactions, formerly unknown pathways, protein complexes, and/or functional modules related to a disease(30, 31). Susceptibility genes for complex diseases tend to be clustered in close network proximity to other susceptibility genes in the protein-protein interactome(32). Therefore, one way to characterize the potential functional impact of a GWAS gene is to comprehensively identify and symbolically represent its interactions with other proteins in a protein-protein interaction (PPI) network. In addition, finding shorter network paths induced via newly identified interactions may help to explain disease phenotypes, especially if the network edges represent physical interactions.

In this study, we experimentally identified protein-protein interactions of HHIP via affinity purification mass spectrometry (AP-MS) in two relevant lung cell lines, lung fibroblasts (IMR90) and bronchial epithelial cells (16HBE). We then combined these experimental interactions with a PPI network derived from publicly available protein-protein interaction databases. We defined HUBRIS as a PPI network composed of interactions in which the products of gene expression (proteins) physically bind each other (i.e., direct PPIs). Moreover, we filtered the proteins represented in our network based on cell-type specific gene expression, giving rise to cell-type specific PPI networks in which the nodes can be associated directly with genes. We then validated several experimentally identified PPIs with co-precipitation assays and investigated the functional implications of newly identified protein interactions with HHIP. We hypothesized that assessment of HHIP protein-protein interactions in relevant lung cell types would identify functional interactions between COPD GWAS genes and suggest new biological mechanisms for HHIP’s role in COPD pathogenesis.

## Results

### AP-MS reveals new interactors of HHIP

We utilized 16HBE (human bronchial epithelial cells, immortalized via SV40 LT antigen) and IMR90 (fibroblasts isolated from human fetal lungs, immortalized by hTERT) cell lines to identify protein interactors of HHIP that may play physiological roles in COPD. We used mass spectrometry analyses of cellular lysates to assess intracellular protein-protein interactions. We performed triplicate assays in both cell lines infected with either an HA-tagged human HHIP lentiviral construct or an empty vector that were subsequently selected with blasticidin to eliminate non-infected cells.

We pulled down HHIP with HA antibody to the HA-tag and found binding of multiple proteins in comparison to the empty vector controls in IMR90 and 16HBE cells. The IMR90 results showed much less non-specific binding to empty control vectors. Therefore, we repeated the 16HBE experiment and analyzed the results of these two 16HBE experiments together. Notably, endogenous HHIP is expressed in IMR90, in contrast to 16HBE cells, which might in part explain the experimental differences between IMR90 and 16HBE cell lines.

We used two analytical approaches to identify protein-protein interactions: Genoppi(33) and SAINTexpress(34). A comparison of results from these two methods is shown in Supplementary Figure *S1*. Notably, Genoppi generally identifies more interactions than SAINTexpress. After examination of the peptide spectral counts (PSC), we opted to focus on SAINTexpress results, which appeared to demonstrate clearer support for significant interactions in the underlying PSC distributions.

Seventy-seven significant HHIP interactors were identified in IMR90 and 34 significant interactors were identified in 16HBE. However, only three of these interacting proteins overlap between these two cell lines (UGGT1, TUBB4B, and HSPA5), suggesting substantial cell-type specificity in protein-protein interactions. TUBB4B and UGGT1 were identified in all three experiments. While all of the significant interacting proteins had good support in the underlying PSC distribution in both pulldown and matched control experiments, we stratified our results according to whether the interactors ever appeared in other mass spectrometry control experiments represented in the CRAPome database(35), which may represent false-positive interactions. After augmenting with CRAPome controls in the analysis, 10 and 18 significant interactions for HHIP remained in IMR90 and 16HBE cells, respectively; however, the overlap between the interactions in both cell lines increased from 3 to 5. For network building, our primary analyses used only the CRAPome-filtered interactions with HHIP, but we performed secondary analyses using all the significant interactions identified in our controlled experiments. A summary of the significant interactions identified with SAINTexpress (with and without CRAPome controls) is provided in Supplementary Table S1. Furthermore, functional enrichment analyses of all the different strata of significant links (with and without CRAPome controls, both cell lines separately, and together) are shown in Supplementary Table *S2*.

Thus, our initial results identified multiple novel protein interactions with HHIP and support the notion that there are likely cell type-specific protein interactions among the COPD GWAS gene products. A visualization of the newly identified cell-type specific interactions of HHIP, after CRAPome filtering, is shown in Figure 1. Additionally, the network of HHIP interactions validated in SAINTexpress (without the CRAPome controls) is shown in Supplementary Figure S2.

**Figure 1:**
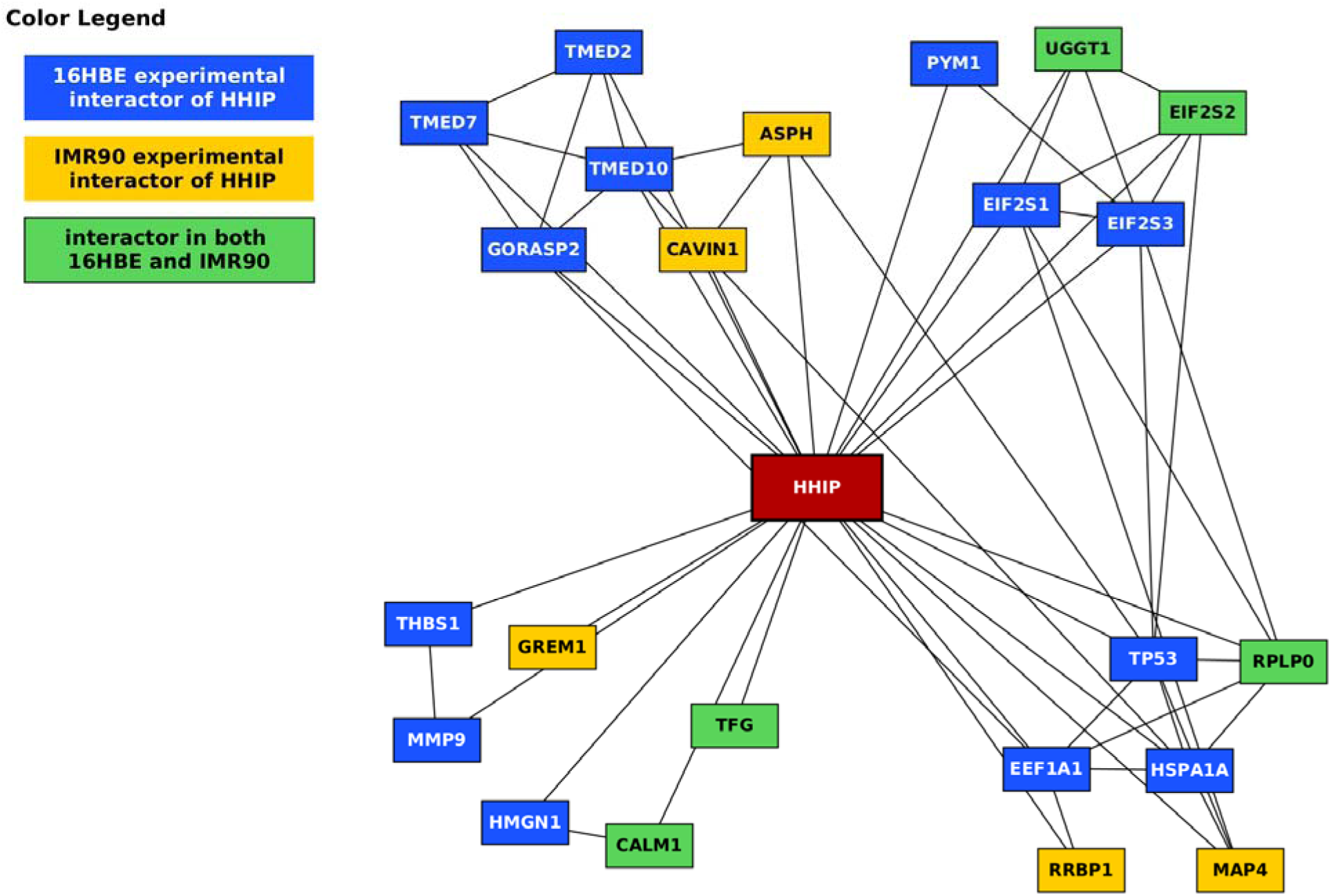
The experimental protein-protein interactions found for HHIP using SAINTexpress and CRAPome validation pipeline. The inter-links between the nodes other than the ones to HHIP are from the HUBRIS network (introduced in the next section). The small clusters emerge due to a force-directed layout algorithm (connected nodes attract; disconnected nodes repel). The clusters suggest denser local connections as well as shared functional roles. Apart from HHIP, all nodes are expressed in their respective cell lines according to our RNA-Seq filtering criteria. The network of HHIP interactions validated in SAINTexpress (without the CRAPome controls) is shown in Supplementary Figure S2.

### HUBRIS reveals the network neighborhood of HHIP in public protein-protein interaction databases

To complement our experimentally determined protein-protein interactions with HHIP, we examined the interactions of HHIP in the literature and in several publicly available databases. To conduct a thorough review, we constructed a large protein-protein interaction (PPI) network by merging 8 PPI databases, namely: HIPPIE(36), HumanNet(37), HURI(38), BioGrid(39), Reactome(40), Interactome3D(41), NCBI(42), and StringDB(43) (in StringDB we only included edges with a non-zero experimental score). We named the emergent network HUBRIS based on a rough acronym of the names of the constituent databases. The merging procedure is described in detail in the Methods section.

Our goal with HUBRIS was not to create a novel pipeline of validating interactions or a new consensus database, but to create a comprehensive picture of the already existing, experimentally determined interactions. Nonetheless, we performed some filtering in the network; for example, we only use edges that can be found in at least two of the constituent databases. Our rationale is that if an interaction passes at least two of the complex curation pipelines of the above-mentioned databases, it is stronger evidence that it likely exists. It is worth noting that many of the PPI curation pipelines are not independent of each other; however, each database uses a different pipeline, combining both automated and manual curation methods.

To better understand the cell type-specific network relationships between HHIP and other proteins, we need to consider that not all proteins in the PPI network are present in a specific cell type. To this end, we used RNA-Seq data obtained from the two cell types used in this study, IMR90 and 16HBE, as a proxy for protein expression to filter nodes in the HUBRIS network (For more details, see Methods). The cell line-specific versions of HUBRIS were derived by removing the respective set of unexpressed nodes from the network and keeping the largest connected component (LCC). The HUBRIS network (without cell type-specific filtering) consists of N=20,103 nodes and E=896,648 edges. The IMR90 and 16HBE specific HUBRIS networks consist of N=11,988, E=619,857 and N=12,226, E=634,957, nodes and edges, respectively. The workflow of generating the cell type-specific HUBRIS networks is shown in Figure 2.

**Figure 2:**
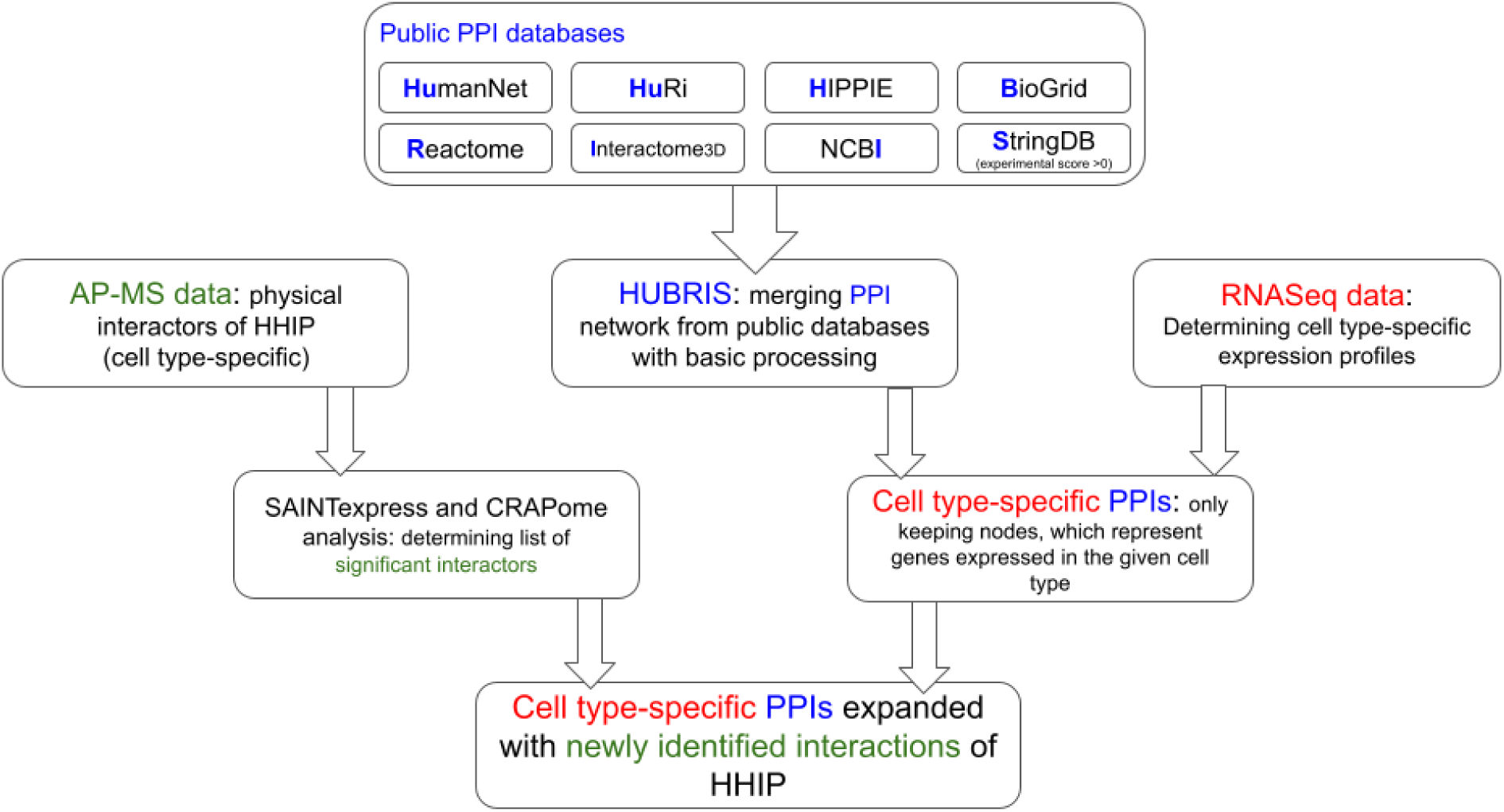
HUBRIS creation pipeline. Each node in this pipeline represents a different step in the process of creating the final, cell type-specific version of the HUBRIS network, with and without the experimental protein-protein interactions from our mass spectrometry analysis. Each step of this process is described in a separate section in Methods. The colors highlight the different strands of data processed and utilized to create the final network versions.

Our HUBRIS network contained the canonical hedgehog pathway interactions with HHIP, with the addition of several other proteins, as shown in Figure 3. In Figure 3, we highlight the proteins that are expressed in our cell lines as well as some details about the experiments in which the interaction was found.

**Figure 3:**
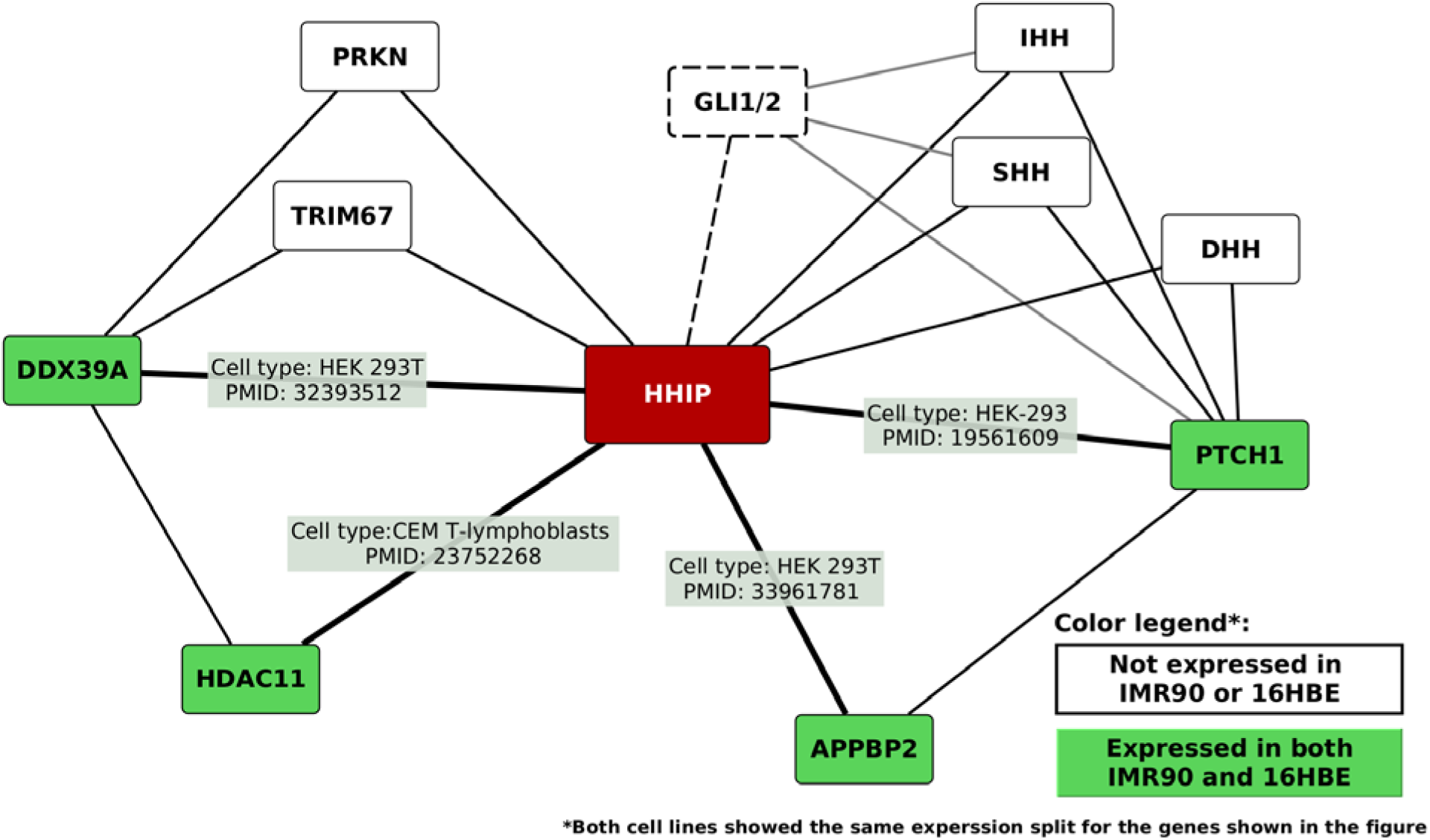
The protein interaction network of HHIP in HUBRIS. Green nodes are expressed in both IMR90 and 16HBE cells, while white nodes are not. On each edge between HHIP and expressed (green) proteins we highlight the cell type, method and PMID of the study in which the interaction was identified (extracted manually from the constituent databases). We used a dashed line for the GLI1/2 interaction because it is not in our HUBRIS database, but it can be found in the OTAR and Wang and Loscalzo et al. databases.

Moreover, we also examined other consensus databases such as OTAR(44), which parses a different set of PPI databases (the only overlap being Reactome) and a consensus database provided by Wang and Loscalzo(45), to further validate our results. We found that both consensus networks confirmed a subset of the interactions found by HUBRIS (the canonical interactions within the HH pathway) and showed an additional interaction with GLI1/2. However, this interaction is gene regulatory rather than protein-protein binding(46).

### Newly identified HHIP interactions reveal shorter paths to other COPD GWAS gene products

Multiple genes have been found to be associated with COPD through GWAS and subsequent functional studies. However, the relationships between these genes in COPD pathogenesis are largely unknown. Using network analysis, our goal is to elucidate whether some of the COPD susceptibility genes have related functional roles in COPD pathogenesis by trying to find paths that show their network proximity. To achieve this, we used HUBRIS together with our newly uncovered experimental interactions and examined the shortest paths between HHIP and a set of COPD GWAS gene products.

We conducted a shortest path analysis (see Methods) separately in HUBRIS networks filtered based on IMR90 and 16HBE RNA-Seq data. We determined all shortest paths between HHIP and the other COPD GWAS gene products in the cell type-specific HUBRIS networks both before and after introducing our newly identified experimental interactions. For a comprehensive summary of how the length and number of shortest paths to the COPD GWAS change after the addition of the new edges see Supplementary Figure S3.

Notably, we found that with the introduction of our new experimentally determined edges along with public PPI data, there are new, *shorter* paths between HHIP and desmoplakin (DSP), a COPD GWAS gene(47) product, and an important component of desmosomes for cellular junctions, in both the IMR90 and 16HBE networks. However, the proteins connecting DSP to HHIP are different in the IMR90 and 16HBE networks. In 16HBE, one of these bridging proteins is TP53. Interestingly, the same three neighbors that form a new path to DSP also connect HHIP to another COPD GWAS gene, TET2 (TP53, EEF1A1, and HSPA1A). It has been recently found that inactivation of *Tet2* in mouse hematopoietic cells increased the risk of cigarette smoke-induced emphysema and inflammation(20). In the IMR90 network there is a new path of length 2 between HHIP and FBLN5, an extracellular matrix component linked with the adhesion of endothelial cells, which is both near a COPD GWAS locus(48) and a cause of cutis laxa(49), a Mendelian syndrome which can include emphysema in its syndrome constellation. In 16HBE cells, we found a number of new shortest paths of length 3 to MFAP2, yet another GWAS gene associated with connective tissue organization. There are no overlapping new shorter paths between IMR90 and 16HBE. We show the emergent network of these paths in Figure 4. The networks emerging when using the SAINTexpress validated edges (no CRAPome controls) are shown in Supplementary Figures S4 (IMR90) and S5 (16HBE).

**Figure 4:**
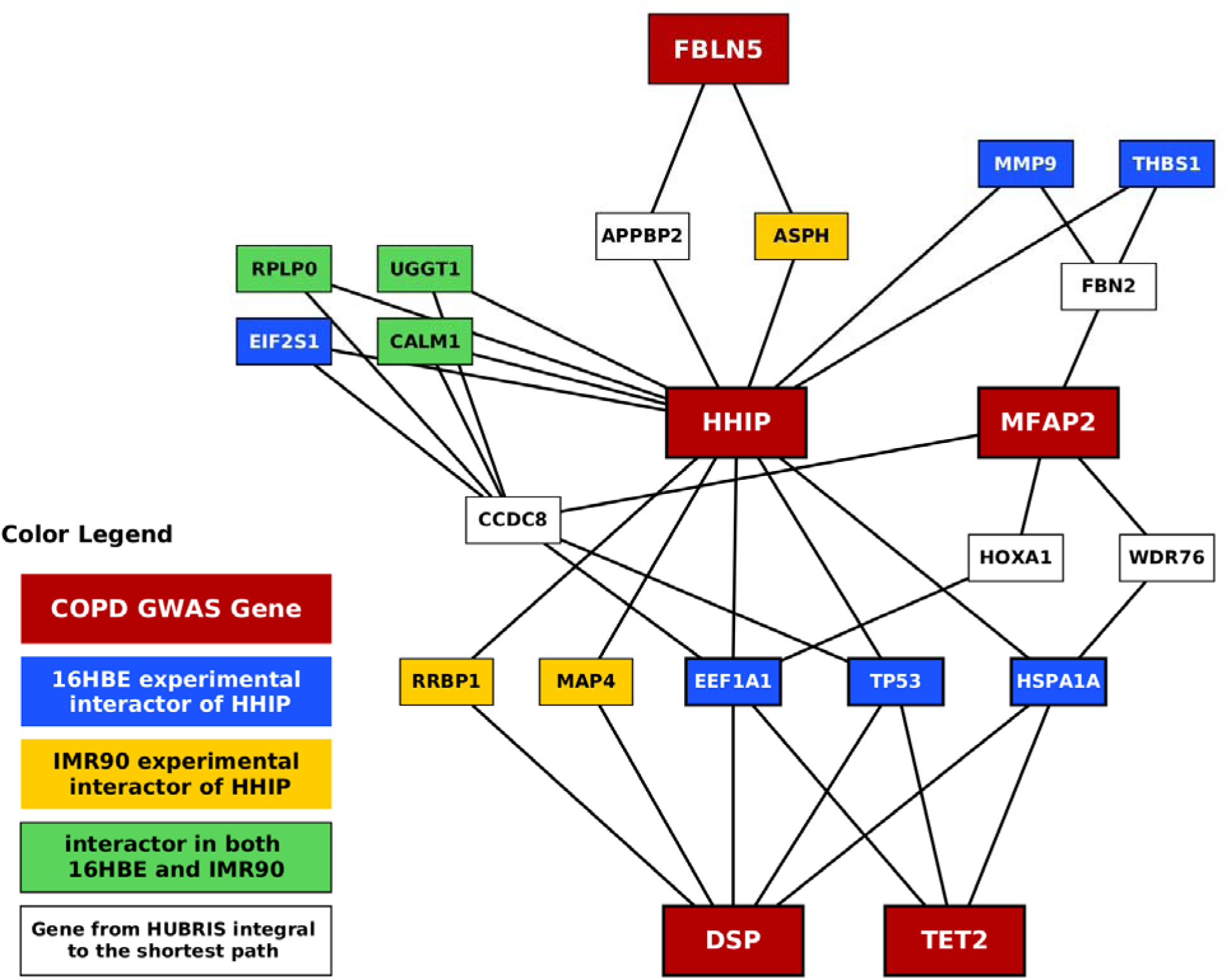
New, shorter paths between HHIP, DSP, MFAP2, TET2 and FBLN5 created with experimental PPI information. The two cell lines revealed different interactors that connect HHIP to the COPD GWAS genes. The different colors highlight the interactors discovered in different cell lines. The networks emerging when using the SAINTexpress validated edges (no CRAPome controls) are shown in Supplementary Figures S4 (IMR90) and S5 (16HBE).

### New shortest paths between COPD GWAS gene products are enriched for proteins involved in extracellular and tissue organization

As noted above, the addition of experimental protein-protein interaction edges from IMR90 and 16HBE cells into the cell type-specific HUBRIS networks revealed new shortest paths connecting HHIP to several COPD GWAS gene products. In unweighted, undirected networks there are often many equivalent shortest paths between two nodes. This means that these new paths are not shorter than the paths already found in the HUBRIS network (the paths found to be shorter are discussed in the previous section), but offer alternative, equivalent pathways to the COPD GWAS gene products. We created a gene set for both cell lines only including genes that are *part of new shortest paths*, emerging via the new experimental interactions. For the details of how the new shortest path genes are found, see Methods.

We then performed functional enrichment analysis on these sets of genes. This revealed that in IMR90 cells these new shortest paths are enriched for genes in the PI3K-Akt pathway (KEGG pathways 2021:FDR=1.02e-10), focal adhesion (KEGG pathways 2021: FDR=5.36e-06, GO:0005925: FDR=1.15e-09), adherens junction (KEGG_2021:FDR=1.18e-05), tight junction (KEGG pathways 2021:FDR=9.61e-05), actin cytoskeleton regulation/reorganization (KEGG pathways 2021:FDR=1.83e-07, GO:0015629: FDR=2.6e-04, GO:0031532: FDR=0.027), stress fiber (GO:0001725: FDR=0.007), Hippo pathway (KEGG pathways 2021:FDR=4.9e-04), and TGF-beta signaling (KEGG pathways 2021:FDR=2.3e-03) For the top 20 list of enriched pathways in IMR90 cells, see Supplementary Figure S6 and for the full results table see Supplementary Table S3.

In 16HBE cells, the new paths reveal enrichments for the PI3K-Akt pathway (FDR=1.01e-13), TGF-beta signaling (KEGG pathways 2021:FDR=8.67e-06), focal adhesion (KEGG pathways 2021: FDR=1.69e-05, GO:0005925: FDR=1.21e-07), adherens junction (KEGG pathways 2021:FDR=1.79e-07), regulation of actin cytoskeleton (KEGG pathways 2021:FDR=9.44e-07, GO:0031532: FDR=0.038), tight junction (KEGG pathways 2021:FDR=2.01e-04), and Hippo pathway (KEGG pathways 2021:FDR=1.88e-04). However, we observe more enrichment for cancer-related pathways than in the case of IMR90 cells, likely because of the direct connection of HHIP to TP53 in 16HBE cells. The top 20 list of enriched pathways in 16HBE cells are shown in Supplementary Figure S7 and for the full results table see Supplementary Table S3.

We highlight the pathways above from the full pathway enrichment analysis because they are critical in lung tissue organization, especially for epithelial cells. The damage, destruction and faulty healing of the lung epithelium is likely a critical determinant of COPD; thus, the emergence of related pathways in the analyzed PPI networks points to better mechanistic explanations as well as potential drug targets.

To reveal all *potential* paths between COPD GWAS proteins independent of cell type-specific gene expression, we also examined a more permissive network setting where we included experimentally identified links from both IMR90 and 16HBE experiments (with CRAPome filtering) and looked for shortest paths in an unfiltered network (i.e., not filtered based on RNA-Seq expression). The paths revealed by the new experimental edges show even more striking enrichment for tissue organization via pathways such as ECM-receptor interaction (KEGG pathways 2021: FDR=7.77e-08) and TGF-beta signaling (KEGG_2021: FDR= 3.62e-03). In the Gene Ontology (GO) Biological Processes pathway set (2021), the most significantly enriched pathways were extracellular structure organization (GO:0043062: FDR=1.40e-21), external encapsulating structure organization (GO:0045229: FDR=1.40e-21), extracellular matrix organization (GO:0030198: FDR=3.46e-20), and collagen fibril organization (GO:0030199, FDR=8.71e-19) as well as some of the pathways highlighted in the cell type-specific cases (Supplementary Figure S8). The specific genes that are common in many of these pathways are related to collagen regulation such as COL1A1; COL1A2; COL2A1; COL4A2; etc. This is due to paths created through MMP9 and THBS1, which are significant interactors of HHIP in 16HBE cells, to COPD GWAS genes MMP1 and MMP12 through a cluster of collagen-regulating genes (see Supplementary Figure S9). The reason that this is not immediately apparent in the different cell line-specific networks is because MMP1 and MMP12 are not expressed in 16HBE.

We also conducted functional enrichment analysis on the nodes in *all* shortest paths between HHIP and the other COPD GWAS genes. Pathways such as the *PI3K-Akt signaling pathway*, *TGF-beta signaling pathway* and *Focal adhesion* remain significant with the inclusion of all shortest paths (nodes of the induced graph between HHIP and the COPD GWAS genes). Moreover, the same pathways are also significant without CRAPome filtering. The resulting significant pathways are listed in Supplementary Table S4.

### Validation and Functional Assessment of Selected HHIP Protein-Protein Interactions

Based on previous studies of COPD pathogenesis, we selected three newly identified experimental interactors of HHIP from our mass spectrometry experiments and network analysis for validation studies: TP53, CAVIN1, and MMP9. As shown in Figure 5, the mass spectrometry interactions were confirmed with co-immunoprecipitation assays for CAVIN1 (IMR90) and TP53 (16HBE), but not for MMP9 (Supplementary Figure S10).

**Figure 5:**
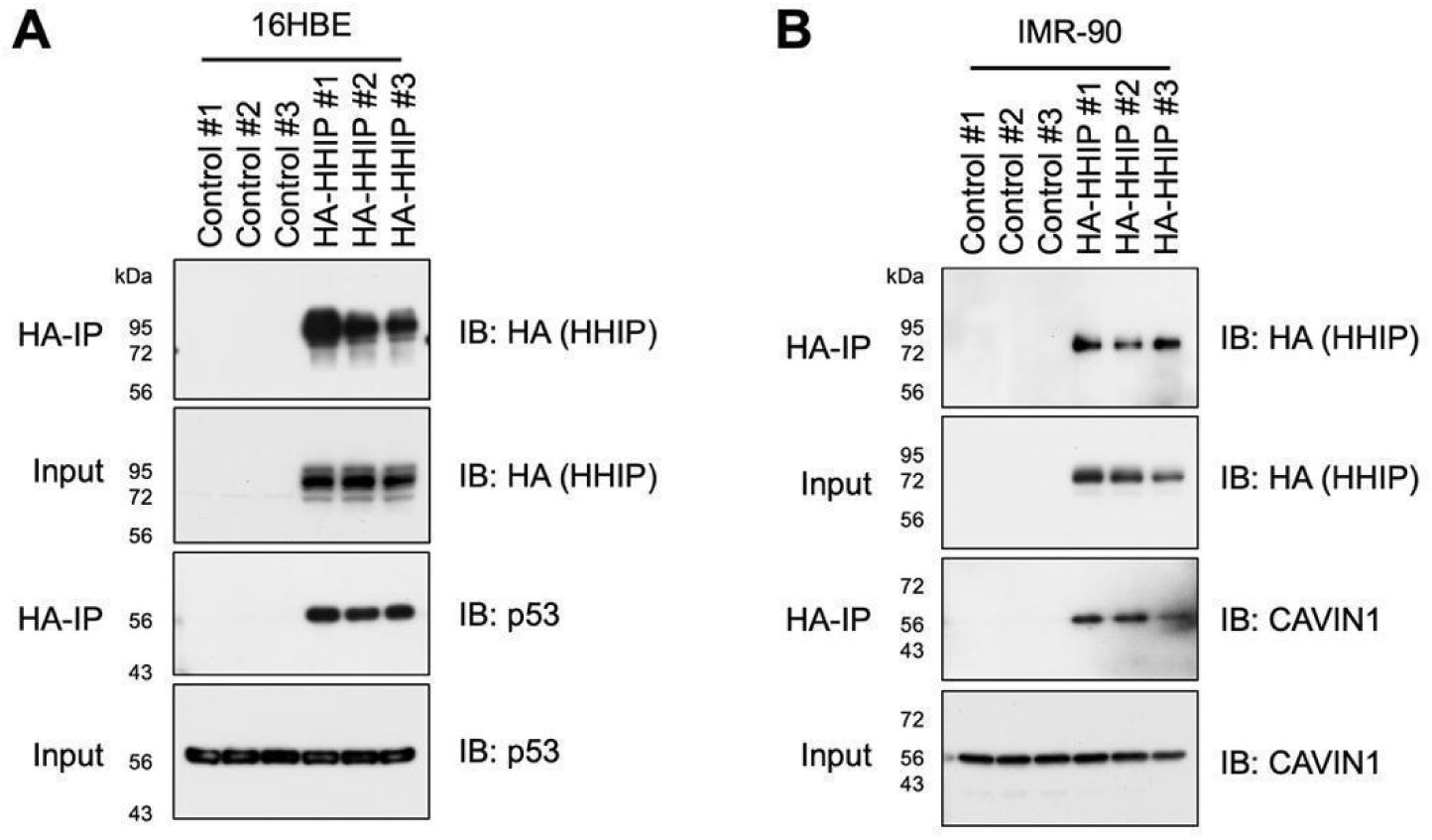
Experimental validation of HHIP-interacting proteins in 16HBE and IMR-90 cells. A-B. Immunoblot (IB) analysis of triplicates of eluted HA-immunoprecipitates (IPs) and inputs derived from 16HBE (A) for TP53 (also referenced as p53) or IMR90 cells (B) for CAVIN1 stably expressing HA-HHIP.

To investigate the biological connections between HHIP and TP53, we reanalyzed a previously performed single cell RNA-Seq analysis of lung tissue from *Hhip^+/-^*and *Hhip^+/+^* mice(50). As shown in Figure 6, P53 (TP53) pathway score was significantly elevated in mouse lung ciliated cells from *Hhip^+/-^* mice. This analysis complements our protein-protein interaction results and suggests that HHIP is likely involved in P53-related functions such as aging and/or cellular senescence.

**Figure 6:**
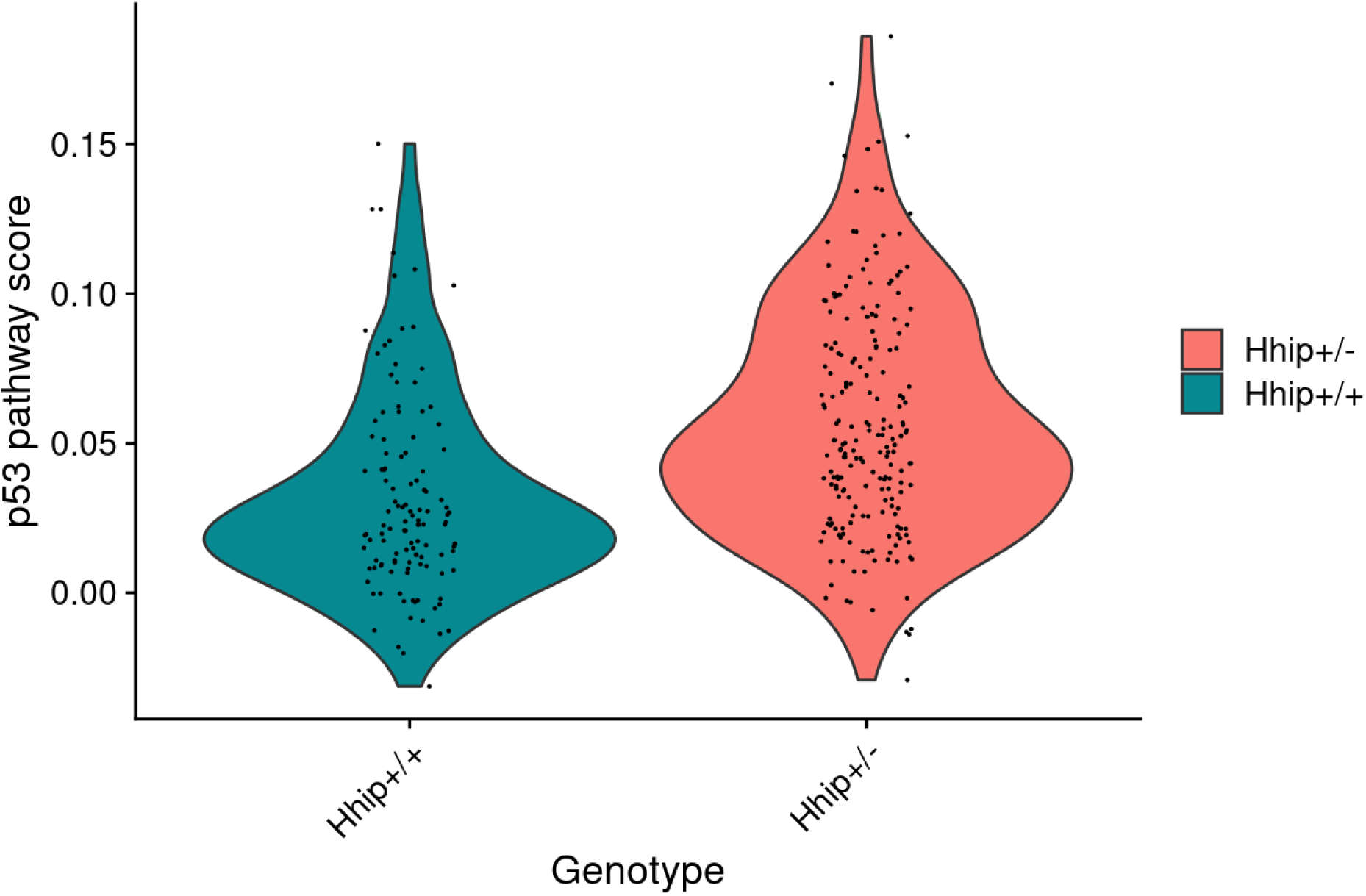
Single cell RNA-Seq was performed in 10 Hhip^+/-^ and 8 Hhip^+/+^ mice. P53 (TP53) pathway scores from the MsigDB Hallmark pathway were compared in ciliated cells. The pathway scores were significantly different at p<6E-10 based on the Wilcoxon test.

Like HHIP-TP53, an interaction between HHIP and CAVIN1 has not been previously reported. To investigate the potential impact of CAVIN1 on COPD pathogenesis, we used siRNA to knock down *CAVIN1* in IMR90 cells and measured RNA levels for a variety of inflammation and senescence-related biomarkers. As shown in Figure 7, knockdown of *CAVIN1* led to a significant increase in the expression of IL6, a key inflammatory cytokine, but did not have a significant effect on IL8, IL33, SIRT1, or SIRT3.

**Figure 7:**
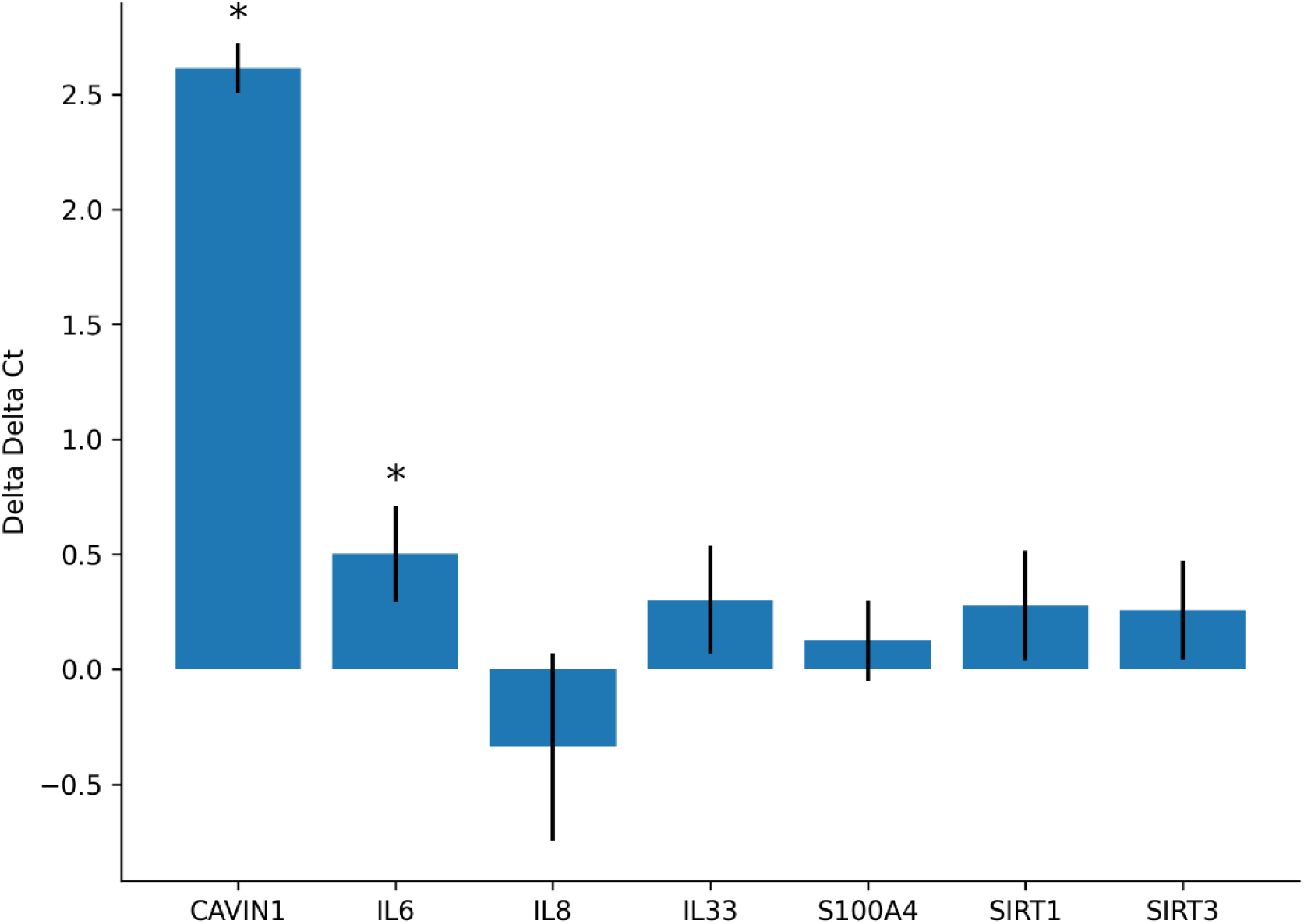
Expression of key genes after CAVIN1 knockdown in IMR90 cells. CAVIN1 siRNA was applied to IMR90 cells, and gene expression (delta delta Ct value) was assessed for several cytokines and cellular senescence markers. The effects of siRNA were assessed using multiple linear regression. *p<0.05; **p<0.01 (Means+/-SD were shown from more than three independent biological replicates)

## Discussion

In this paper, we have explored the protein-protein interactions of HHIP, a well-established COPD GWAS gene, in two different lung cell lines. We used AP-MS experiments to determine cell type-specific interactions of HHIP. We also built a large PPI network consisting of 8 different public databases (HUBRIS). Using RNA-Seq data to filter out genes that are not expressed in IMR90 or 16HBE cell lines, we created cell type-specific versions of HUBRIS. We used these networks to examine the network neighborhood of HHIP in cell type-specific settings, as well as the different network paths connecting HHIP to other COPD GWAS gene products. The newly discovered interactions showed new *shorter* paths to some COPD GWAS gene products and new alternative paths, which are enriched for cellular functions such as extracellular matrix and tissue organization. We also validated two novel HHIP protein interactions (with CAVIN1 and TP53) by co-immunoprecipitation and performed functional studies of these validated interactions that provide insights into the potential mechanisms by which HHIP influences COPD genetic risk.

Recently, Pintacuda et al.(51) and Hsu et al.(52) demonstrated that mapping experimental, cell type-specific protein-protein interaction networks for GWAS gene products has great potential for understanding complex disease pathobiology. Their experiments were conducted in brain cells, mapping connections between (among others) GWAS genes associated with neurological disorders and found that the interactomes were enriched for genes related to the pathogenesis of the respective diseases. These studies showcase that finding cell type-specific protein-protein interactions of disease-related genes can be an important step in understanding complex disease pathogenesis.

Two validated protein interactions of HHIP show particularly promising functional ties to COPD pathogenesis: CAVIN1 and TP53. CAVIN1 was recently identified as a protein biomarker of COPD in COPD lung tissue(53), and it has also been implicated in COPD pathogenesis based on correlation-based network analysis of transcriptomic data in COPD vs. control lung tissue(54). A recent study has shown that CAVIN1 can regulate cellular responses to oxidative stress by inhibiting NRF2, a key transcription factor of antioxidant genes that regulates oxidative stress(55). Reactive oxygen species (ROS) can release CAVIN1 from caveolae through lipid peroxidation, which in turn inhibits NRF2 and thus promotes apoptosis (by reducing the antioxidant response). Such regulation of the stress response of cells creates a delicate collective balance, which determines tissue health, integrity, and healing. By showing a direct protein interaction between HHIP and CAVIN1, we have identified one possible explanation for the way genetic variants that regulate the expression of HHIP could influence tissue integrity and response to oxidative stress. Moreover, ferroptosis, a specific cell-death program associated with lipid peroxidation, NRF2 regulation, and iron metabolism, could be linked to another COPD GWAS gene product, IREB2, whose aberrant behavior causes abnormal levels of iron in the cytosol and in the mitochondria(6).

Previously, in *Hhip*^+/-^ mice that demonstrated more severe airspace enlargement after chronic cigarette smoke exposure or aging than wild-type mice, we detected increased levels of p53 (TP53) in their lungs, suggesting potential regulation by HHIP of p53 signaling(27). We have now demonstrated that HHIP and p53 have a direct protein interaction. We also reanalyzed murine single cell RNA-Seq data from *Hhip^+/-^* and wild-type mice, demonstrating increased P53 pathway activity in lung ciliated cells. Primarily, p53 is known as a key tumor suppressor, regulating a number of cell-cycle checkpoints, DNA-damage repair, cellular senescence, and apoptosis. However, p53 also plays a role in directly maintaining redox homeostasis. For instance, p53 can be activated by AMPK, a protein which responds to lowered ATP levels in mitochondria, a condition often caused by oxidative damage(56). P53 prevents apoptosis but can downregulate Parkin and DRP1, which are responsible for mitophagy and mitochondrial fission, respectively, thus contributing to increased ROS(57). p53 can also induce the transcription of several antioxidant enzymes (such as SOD2 and GPX1)(58). On the other hand, in extended oxidative stress p53 switches its role and promotes apoptosis by inhibiting antioxidant transcription factors such as NRF2(59) as well as anti-apoptotic proteins such as BCL-2(60). Both excessive cell death and cellular senescence under sustained damage could potentially lead to COPD. We have shown in this paper that HHIP interacts with several key proteins that regulate stress response and make important decisions determining the fate of cells. Further study into the nature of these interactions may reveal causal mechanisms which lead to the pathogenesis of COPD and related diseases.

These network analysis results also confirm some previous insights from *HHIP* knockdown experiments, where the network modeling revealed the most significant effect of the knock-down affected genes related to ECM and cell growth and proliferation regulation(61). The composition of the ECM is key in maintaining the required conditions for healthy gas exchange in the lungs. Numerous studies show that the ECM composition changes during the development of COPD(4, 62, 63). The pathways through MMP9 and Thrombospondin 1 (THBS1) to Fibrillin (FBN2) and MFAP2 are all directly associated with the regulation of elastic fibers, collagen, and other essential ECM components responsible for the physical properties of lung tissue. These new links have the potential to reveal mechanisms by which HHIP directly influences airway mechanics.

Our study has several limitations. First, there is very little overlap between our set of experimental protein-protein interactions and the set of HHIP interactions found in the literature.

This could be because the other experiments were conducted in other cell types (see Figure 3), and that interactions in public PPI databases are not exclusively from AP-MS experiments (nor are they exclusively protein-protein physical interactions). For example, in StringDB the “experiments” channel, which scores edges with experimental evidence, may include relationships beyond direct protein-protein physical binding experiments, such as co-expression after CRISPR knockdown. Additionally, each AP-MS experiment only captures a fraction of all the possible interactions; although we performed our experiments in triplicate, detection of protein-protein interactions was likely not complete(64). The lack of overlap can also be attributed to the limited composition of existing PPI databases, as indicated by the observation that there are many different alternative databases with often very little overlap between them (see Supplementary Figure S11). This poses some additional limitations on our own HUBRIS network and the subsequent network analysis, namely that there are multiple levels of uncertainty regarding the existence of interactions (or lack thereof). One of the specific goals of our study is to address this limitation by augmenting the PPI network with high quality experimental interactions in disease-relevant cell types. Another limitation is that the mass spectrometry experiments may not be physiological, as HHIP is not expressed in 16HBE cells. However, HHIP is highly expressed in a subset of human alveolar type 2 epithelial cells(65) and can also be detected in human primary bronchial epithelial cells (based on data from (66)), suggesting that the protein-protein interactions that we identified in 16HBE cells are biologically relevant. The relationship between the interactions that we identified in 16HBE cells with non-physiological expression of HHIP and the endogenous interactions in alveolar type 2 epithelial cells requires further study. Moreover, our failure to validate the MMP9 - HHIP interaction with co-immunoprecipitation indicates that the detected interactions may not be all valid, or they may be indirect. Finally, for the creation of the cell type-specific networks, we used RNA-Seq data for the filtering of nodes instead of proteomics. This can cause inaccuracies in the network analysis as some proteins may exist in a cell but no longer have measurable RNA (false negatives), or proteins can be degraded post-translationally (false positives).

Despite these limitations, our results suggest that the identification of protein-protein interactions in relevant cell types can provide important insights into the biological functions of complex disease GWAS genes.

## Methods

### Experimental Materials and Methods

#### Cell culture

Human bronchial epithelial 16HBE and embryonic kidney epithelial 293T cells were purchased from ATCC (Minnesota, VA). Human telomerase reverse transcriptase (hTERT)-immortalized human fetal lung IMR-90 cells were obtained from Dr. William Hahn’s laboratory (Dana-Farber Cancer Institute). 16HBE and IMR90 cells were cultured in Eagle’s minimum essential medium (EMEM). 293T cells were maintained in Dulbecco’s modified Eagle medium (DMEM). EMEM and DMEM were supplemented with 10% fetal bovine serum (FBS), 100 units of penicillin, and 100 μg/mL streptomycin.

#### Lentiviral packaging and infection

Transfection for lentiviral packaging was conducted by the polyethylenimine (PEI) protocol as described previously(67). Briefly, 293T cells were seeded at 3.0×10^6^ cells per 100-mm plate the day before transfection. For transfection, 4 μg of pLenti-CMV-HA-HHIP lentiviral transfer plasmid or pLKO as a negative control, 3 μg of psPAX2, a lentiviral packaging plasmid, and 1 μg of pMD2.G, a lentiviral envelope plasmid, were diluted in 700 μL of serum-free DMEM, and 24 μg of PEI (1 μg/mL) (Polysciences, Warrington, PA) was directly added to the diluted DNA. After vigorous vortexing for 15 sec, the mixture was incubated at room temperature for 15 min and added to cells. After 24 h of incubation, the medium was replaced with fresh medium and collected three times every 24 h. Collected viral-containing media was filtered (0.45 μm) and added to target cells in the presence of 4 μg/mL polybrene. The infection was repeated twice a day. Cells were then selected with blasticidin (5 μg/mL) to eliminate non-infected cells, and HA-HHIP expression was validated by immunoblot analysis.

#### Antibodies and plasmids

Monoclonal anti-HA agarose (A2095), rabbit anti-HA tag antibody (H6908), monoclonal anti-Vinculin antibody (V9131), and HA Peptide (I2149) were purchased from Sigma-Aldrich (St. Louis, MO). Anti-β-Actin antibody (#3700), anti-p53 (7F5) antibody (#2527), anti-Cavin-1 antibody (#69036) and anti-MMP9 (D6O3H) XP Rabbit mAb antibody (#13667) were purchased from Cell Signaling Technology (Danvers, MA). pLenti-CMV-HA-HHIP lentiviral expression plasmid was generated by subcloning the appropriate human HHIP cDNA PCR fragment into pLenti-CMV Blast (659–1) (Addgene). HA-tag was inserted into human HHIP cDNA after Ala 23, next to the signal peptide, using the QuikChange Site-Directed Mutagenesis Kit (Agilent Technologies, Santa Clara, CA) according to the manufacturer’s instructions.

#### Immunoblot analyses

Cells were lysed in NP-40 cell lysis buffer (50 mM Tris-HCl, pH 7.5, 120 mM NaCl, and 0.5% NP-40) supplemented with a protease inhibitor cocktail (cOmplete, Mini Protease Inhibitor Cocktail, Roche, Basel, Switzerland) and phosphatase inhibitors (PhosSTOP, Calbiochem). Protein concentrations of lysates were determined with the Bio-Rad Protein Assay Dye (Bio-Rad Laboratories, Hercules, CA). Forty micrograms of whole-cell lysates were resolved by sodium dodecyl sulfate-polyacrylamide gel electrophoresis (SDS-PAGE) and transferred to polyvinylidene difluoride membranes (Bio-Rad). The membranes were blocked with 5% nonfat dry milk in Tris-buffered saline with 0.05% Tween 20 (TBST, pH 8.0) and probed with antibodies (at 1:1000-1:4000 with 5% BSA in TBST), as shown in Figure 5.

#### Purification of HHIP interacting proteins

For immunoprecipitation, infected IMR90 or 16HBE cells were harvested and lysed in NP-40 cell lysis buffer containing protease and phosphatase inhibitors. Two milliliters (8 mg/mL) of cell lysates were incubated with 40 μL slurry of anti-HA-tag antibody–conjugated beads for 12 h at 4 °C with gentle rocking. The tubes were briefly centrifuged to precipitate the HA-immunoprecipitates (IPs), and the beads were washed five times with 1 mL of NP-40 washing buffer (20 mM Tris, pH 8.0, 100 mM NaCl, 1 mM EDTA, and 0.5% NP-40) and three times with 1 mL of PBS. HA-IPs were eluted with 80 μL HA peptides (0.25 mg/mL) by gentle rocking for 15 min. After repeating the elution three times, 10% of eluates were suspended in 2×SDS sample buffer for immunoblot analysis, and the remaining was subjected to acetone precipitation as described previously(68). To precipitate eluted proteins, the elutes were added four times the sample volume of cold acetone, vortexed, and incubated for 1 h at -20 °C. The samples were pelleted by centrifugation for 10 min at 15,000 rpm, air-dried for 30 min at room temperature, dissolved in 10 mM DTT, and subjected to mass-spectrometry analysis.

#### Tandem Mass spectrometry

The protein pellets were reduced with 10 mM DTT for 30 min, alkylated with 55 mM iodoacetamide for 45 min, and subjected to trypsin/Lys-C digestion overnight at pH=8.0. The resulting peptide mixture was quenched with 10 uL of 5% formic acid and analyzed by microcapillary reversed-phase liquid chromatography-tandem mass spectrometry (LC-MS/MS) using a high-resolution Orbitrap Exploris 480 (Thermo Fisher Scientific, Waltham, MA) mass spectrometer in positive ion DDA mode (Top 12) via higher energy collisional dissociation (HCD) coupled to a Thermo EASY-nLC1200 nano-UHPLC. A 75 µm i.d. x 10 cm microcapillary column packed 3 µm C_18_ beads and a 100 µm i.d. x 3 cm trapping column packed with 3 µm C_18_ beads (ESI Source Solutions, Woburn, MA) were used for LC-MS/MS separation. MS/MS data were searched against the UniProt Human protein database (containing 82,485 entries) using Mascot 2.7 (Matrix Science, London, UK). Data were analyzed using Scaffold Q+S 5 protein identification software (Proteome Software, Portland, OR). Peptides and modified peptides were accepted if they passed a 1% false discovery rate (FDR) threshold.

#### Assessment of HHIP protein interactors

Human reference proteome database (UP000005640) was downloaded from UniProt (accessed on 06/29/2022) to map accession numbers to gene symbols. Only interactors with positive spectral counts in at least two replicates were retained for downstream statistical analysis. We used SAINTexpress and Genoppi software programs to assess significant interactions. For SAINTexpress, significant interactions were defined by an average posterior probability of at least 0.8, Bayesian FDR no greater than 5% and a positive log likelihood ratio. For Genoppi, an FDR cutoff of 5% was used. To further account for potential pull down contaminants, CRAPome database version 1.0 was downloaded and included as additional controls in the SAINTexpress analysis. We kept only single-step AP-MS experiments and removed outliers, leading to a total of 262 additional CRAPome controls.

### Creating HUBRIS

For the creation of the HUBRIS network, we merged 8 different protein-protein interaction databases from the following sources:

- HumanNet: https://staging2.inetbio.org/humannetv3/networks/HS-PI.tsv
- HuRI: http://www.interactome-atlas.org/data/HuRI.tsv
- HIPPIE: http://cbdm-01.zdv.uni-mainz.de/~mschaefer/hippie/hippie_current.txt
- BioGrid: https://downloads.thebiogrid.org/File/BioGRID/Release-Archive/BIOGRID-4.4.219/BIOGRID-ALL-4.4.219.mitab.zip
- Reactome: https://reactome.org/download/current/interactors/reactome.homo_sapiens.interactions.tab-delimited.txt
- Interactome3D: https://interactome3d.irbbarcelona.org/downloadset.php?queryid=human&release=current&path=complete
- NCBI: https://ftp.ncbi.nih.gov/gene/GeneRIF/interactions.gz
- StringDB: https://stringdb-static.org/download/protein.links.detailed.v11.5/9606.protein.links.detailed.v11.5.txt.gz

For all databases the date of download is 01/24/2024. All analyses are based on the state of the databases on this date.

All networks were used in their linked form, except for StringDB, where we performed a preliminary filtering for the experimental score being larger than zero, as we are looking for concrete physical interactions not functional associations. In addition, for databases where non-human species were included, we filtered to include only interactions found in humans.

#### Merging the networks

The first step of the merging required using a common gene/protein identifier as many networks used different identifiers, such as Entrez Gene Id (HumanNet, HIPPIE, BioGrid, NCBI), Uniprot Id (Interactome3D, Reactome), Ensembl Gene Id (HuRI), and Ensembl Peptide Id (StringDB). Since Entrez Gene Id was the most common, we decided to use it as the common id. We then used the BioRosetta tool (https://github.com/reemagit/biorosetta) to re-label the networks *not* using Entrez Gene Id.

In cases where there was no valid or unique translation, we kept the node in the network with the original identifier (as it could still be structurally significant) and appended the shortened form of the *source id type* to the node id with an underscore. For example, the Ensembl Peptide Id ENSP00000266991 did not have a valid translation to Entrez Gene Id in BioRosetta, and therefore the node received the new label ENSP00000266991_ensp. This helps us to manually look up relevant nodes when interpreting results. In the raw, unfiltered version of HUBRIS roughly one-third of the nodes do not have a valid translation in our protocol; however, in the final, filtered version of HUBRIS (N=20,103) there are only 244 such nodes.

After the relabeling of the individual networks, we merged them into a single network that for every edge aggregated the names of the different databases that contained them. We call the number of databases that contain a single specific edge the *database support count* and it is a property of every edge.

#### Properties of the merged HUBRIS

The raw merged network consists of N=32,384 nodes and E=3,233,850 edges. For the edges, the database support distribution is the following: 1: 2,337,191, 4: 304,861, 5: 242,908, 3: 167,029, 2: 158,215, 6: 20,493, 7: 2,946, 8: 207.

#### Filtering the HUBRIS network

The above statistics show that most of the edges (>72%) are from a single database source and these edges are overwhelmingly from StringDB (>89%). Having manually examined parts of the network that are relevant for our study (especially the neighborhood of HHIP and the other COPD GWAS gene products), we decided to trim our network and only keep edges that have support from at least 2 databases. The resulting network has the following statistics: N=32,384 nodes and E=896,659 edges, where the giant component is composed of N=20,103 nodes and E= 896,648 edges.

In all analyses described in the paper and the accompanying documents when we refer to HUBRIS, we refer to the network defined above.

### Cell type-specific filtering of HUBRIS based on RNA-Seq data

To better understand the cell type-specific network processes between HHIP and other proteins, it is necessary to consider that not all proteins present in the PPI network are present in a specific cell type. To account for this effect, we used RNA-Seq data obtained from the two cell types we used in this study: IMR90 and 16HBE.

RNA sequencing was performed from cultured cells by Novogene. Paired end cDNA libraries were constructed and sequenced on Illumina platforms for an average read depth of ∼50 million reads. Reads were aligned to hg37 using the HISAT aligner(69). Gene expression was quantified as Fragments Per Kilobase of transcript per Million mapped reads (FPKM).

For both cell lines, we used the same criteria: we excluded genes with an absolute count of 0 since the normalization can skew these values into nonzero values, and then used a cutoff of 0.25 on the log2 of the FPKM values (see Supplementary Figure S12).

The value of 0.25 was determined to strike an appropriate balance between removing nodes with low expression while still allowing us to retain all proteins measured in the AP-MS experiments (including HHIP).

Finally, we kept genes that were expressed and removed every other node from HUBRIS, and by keeping the remaining largest connected component (LCC) thus created the cell type-specific network.

### Shortest path analysis

The shortest path in a network is defined between two nodes (or vertices) as paths in which the sum of the weights of its constituent edges is minimized. In other words, considering a source node and a target node in an unweighted network, we seek the sequence of edges through which one can reach the target from the source with the minimal number of steps.

HUBRIS (like PPI networks in general) is undirected and unweighted, which typically leads to a large number of equivalent shortest paths between two nodes, especially if the network is locally dense. Since there is no way to differentiate between these sets of shortest paths, we merge them into a sub-network, which we call the ***induced graph***, a graph-theoretical term defined as a subgraph consisting of a subset of the vertices (nodes) of the graph and all the edges (from the original graph) connecting pairs of vertices in that subset.

When analyzing the connections between HHIP and the other COPD GWAS gene products, we generate the induced graph between HHIP and the nodes representing the GWAS gene products. Since no GWAS node is directly connected to HHIP, the subgraph is generated from all equivalent shortest paths and their constituent genes. To study the cell type-specific differences as well as the *new paths* introduced by the edges discovered in our mass spectrometry experiments, we generate four different induced graphs (two cell type-specific networks, both with and without the experimental edges) and compare them.

To find which nodes are part of new shortest paths created by the newly identified experimental edges, we just subtract the set of nodes of the induced graph of the original cell type-specific HUBRIS network from the set of nodes induced from the network expanded with the new edges. An example of a network constructed just from the new paths with the procedure described above (using both IMR90 and 16HBE links and no RNA-Seq filtering) is shown in Supplementary Figure S9.

We conducted the GSEA analysis on the node-sets of different induced graphs as well as on the inferred differences between the expanded vs original networks. For the functional enrichment analysis, we exclude the target set (the COPD GWAS genes) and the set of new neighbors from the analyzed gene set. The results of this analysis are shown in Supplementary Figures S6 to S8 and Supplementary Table S3.

All network analyses were conducted using the NetworkX python package (v 3.2).

Our network analysis pipeline is available on GitHub: https://github.com/deriteidavid/hubris

### Functional enrichment analysis

We conducted a functional enrichment(70, 71) on the different gene sets obtained from the induced networks described in the previous section.

We used the gseapy python package to run the analysis, with the following settings:

- All input node sets were **unranked** (hypergeometric test).
- Gene sets: KEGG_2021_Human, GO_Cellular_Component_2018, GO_Biological_Process_2021
- The background was always set as the node-list of the HUBRIS network.
- In all the figures of the supplement, we ranked the enriched pathways based on the level of significance.
- The significance is FDR corrected by the gseapy package.

The code reproducing our analysis is available on GitHub: https://github.com/deriteidavid/hubris

### Functional Validation Analyses

Single cell RNA-Seq data from the *Hhip* heterozygous and wild-type murine lungs was used to generate the p53 (TP53) pathway score. Briefly, single cell RNA-Seq using 10X Chromium Single Cell 3’ v2 and inDrops for a total of 38,875 cells (26,952 Hhip+/- and 11,815 Hhip+/+ cells) was reanalyzed after quality control, SCTransform and integration(72). AddModuleScore function in Seurat was used to calculate the composite p53 pathway score for 200 genes from Molecular Signature Database Hallmark p53 pathway gene set (https://www.gsea-msigdb.org/gsea/msigdb/human/geneset/HALLMARK_P53_PATHWAY.html).

siRNA knockdown of *CAVIN1* was performed using ON-TARGET plus SMARTpool Human PTRF siRNA (Dharmacon, Cat#L-012807-02-0005) for the experimental group, alongside control siRNA (ON-TARGETplus Non-targeting Control pool, Dharmacon, Cat# D-001810-10-05) for the control group with Lipofectamine RNAimax transfection reagent. Transfection was conducted on 60-70% confluent IMR90 cells using 60 pmol per well of siRNA. Seventy-two hours after treatment RNA was extracted (Zymo Research Quick RNA mini prep kit), followed by RT-qPCR analysis employing gene probes (Supplementary Table S5) to determine expression of target genes.

Statistical analysis of the three biological replicates of the CAVIN1 siRNA experiments was performed using multiple regression analysis using the method described by Yuan and colleagues(73).

## Supporting information

Supplementary Doc

## Author Contributions

D.D. – conceptualization, network generation and analysis, data curation (public databases), software (HUBRIS, network methods, GSEA), writing (original draft), visualization (Figures 1-4, 7, S2-S9, S11, S12)

H.I. – experimental investigation (AP-MS, co-IP), validation, writing (review and editing), visualization (Figures 5, S10)

P.C. – formal analysis (AP-MS data), writing (review and editing), supervision

J.H.Y. – experimental investigation (scRNA-Seq), validation, formal analysis, writing (review and editing), visualization (Figure 6)

Z.X. – formal analysis (AP-MS data), validation, visualization (Figure S1), writing (review and editing

W.J.A. – experimental investigation (siRNA KO), validation, formal analysis, writing (review and editing)

J.M.A – resources (sample analyses), MS data acquisition and analysis, writing (review and editing)

F.G. – experimental investigation (RNA-Seq of cell lines), writing (review and editing)

X. Z. – supervision, writing (review and editing)

K.G. – supervision, writing (review and editing)

W.W. – conceptualization, funding acquisition, supervision, writing (review and editing)

E.K.S. – conceptualization, funding acquisition, project administration, formal analysis (siRNA KO data), supervision, writing (original draft)

