## Supplementary Doc for "HHIP protein interactions in lung cells provide insight into COPD pathogenesis"

### Supplementary Figures

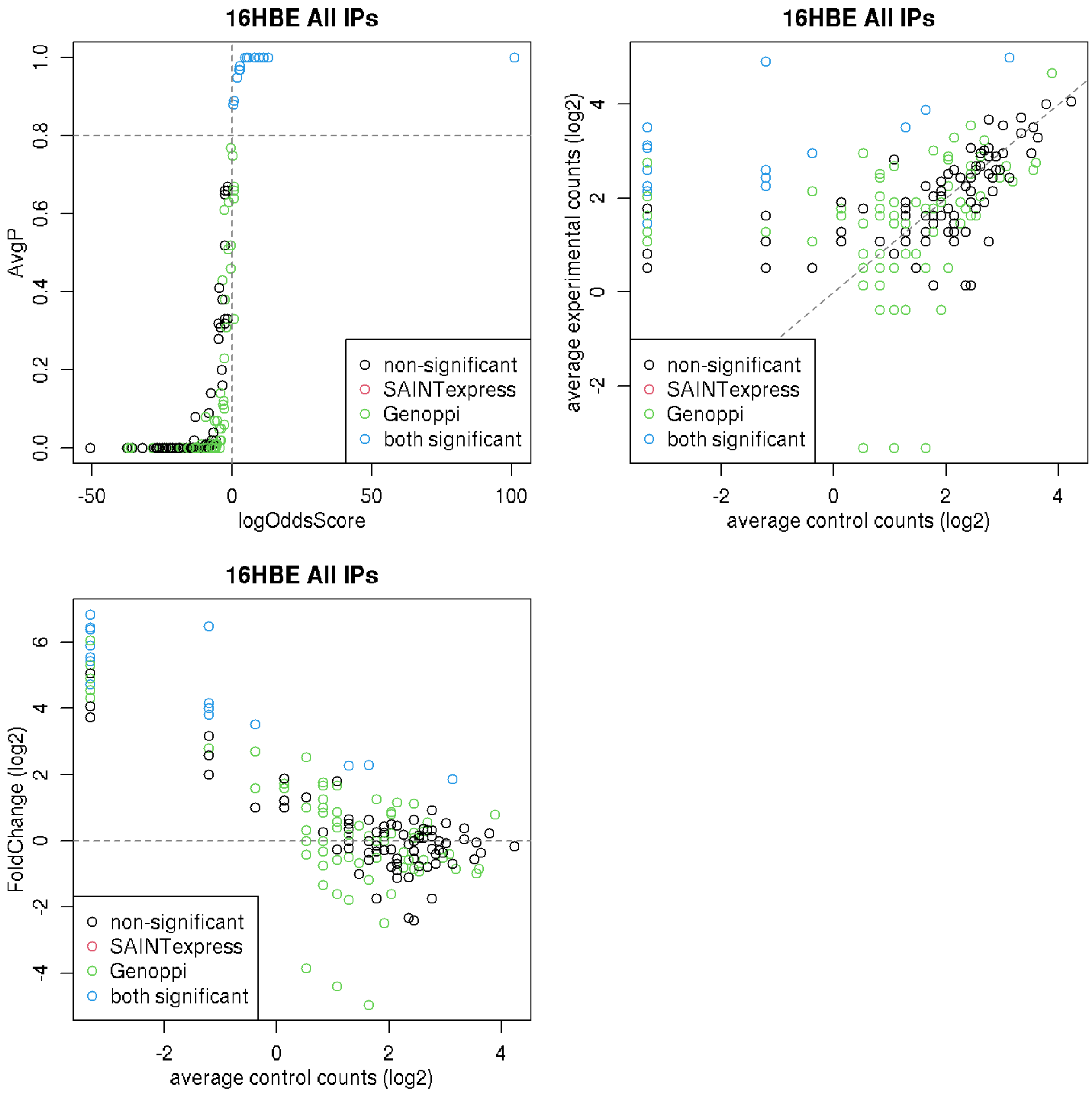


***Figure S1. SAINTexpress and Genoppi comparative analysis on HHIP AP-MS experiments in 16HBE****. Significant HHIP protein interactors identified from the two methods are color coded (black: non-significant, red: significant only in SAINTexpress (no points), green: significant only in Genoppi, and blue: significant in both methods). The top left panel shows the distribution of SAINTexpress statistics (AvgP: average posterior, logOddsScore: log likelihood ratio). The top right panel compares the spectral counts in control and HHIP experiments. The bottom left panel shows the relationship between spectral counts in control experiments and the fold changes between HHIP and control experiments.*

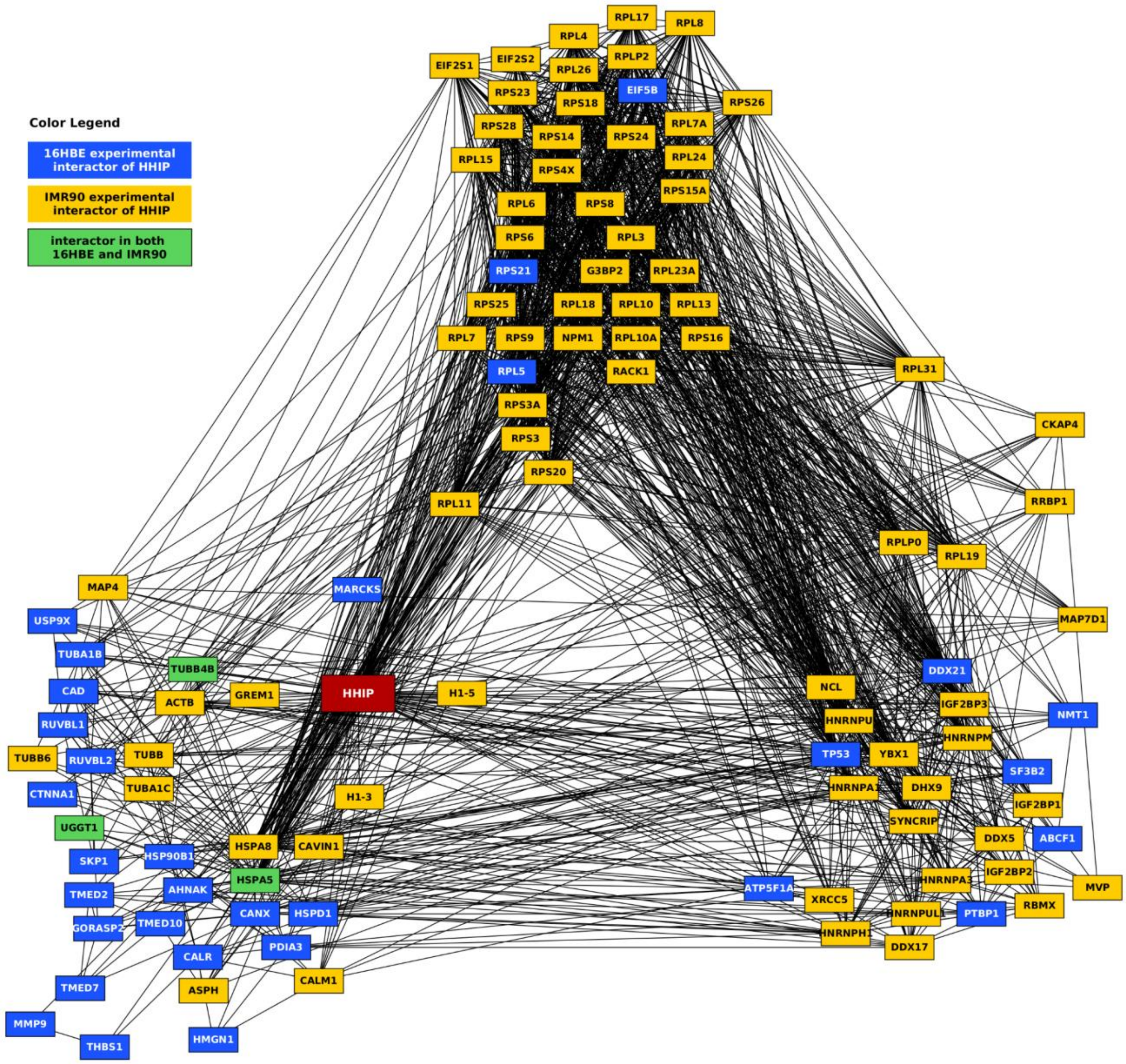


***Figure S2: The experimental interactors of HHIP using the SAINTexpress validation pipeline***. *The experimentally determined protein interactors of HHIP are shown in rectangles. The inter-links between the nodes other than the ones to HHIP are from the HUBRIS network. The clusters are formed due to a force layout algorithm (connected nodes attract; disconnected nodes repel). The clusters suggest denser local connections as well as shared functional roles. With the exception of HHIP, all nodes are expressed in their respective cell lines according to our RNA-Seq filtering criteria. The nodes found in different colors represent the different cell lines (or both).*

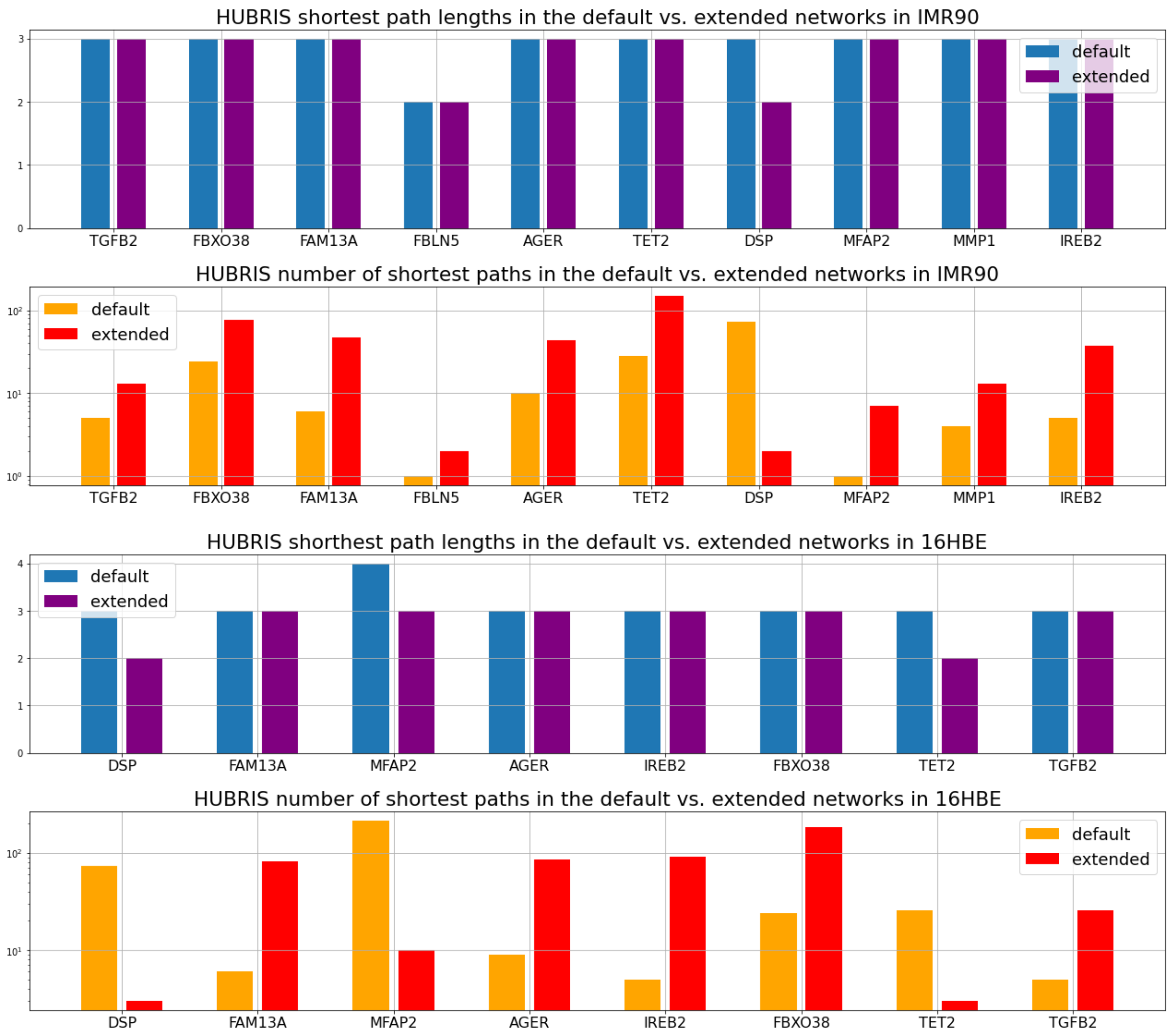


***Figure S3: The length (top) and number (bottom) of shortest paths between HHIP and the selected set of COPD GWAS genes*** *in the default HUBRIS network (only including literature interactions) vs. the extended HUBRIS network, where we also include our newly identified experimental interactions in this study.*

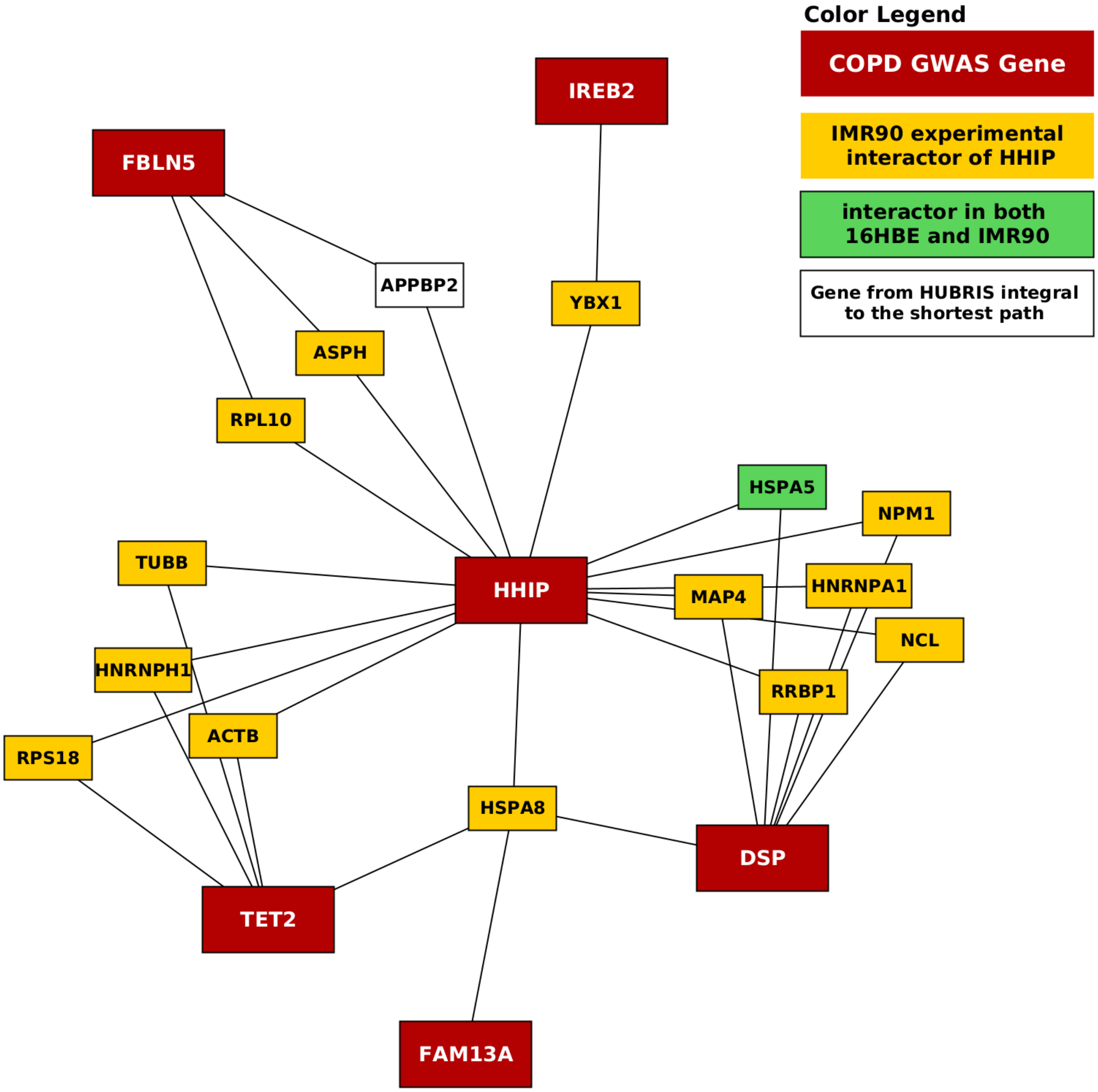


***Figure S4: New, shorter paths between HHIP and COPD GWAS genes in IMR90.*** *The experimental links are determined in SAINTexpress (without CRAPome). The COPD GWAS gene products are larger with a red color, and we also highlight nodes that were parts of shorter paths in 16HBE cells*

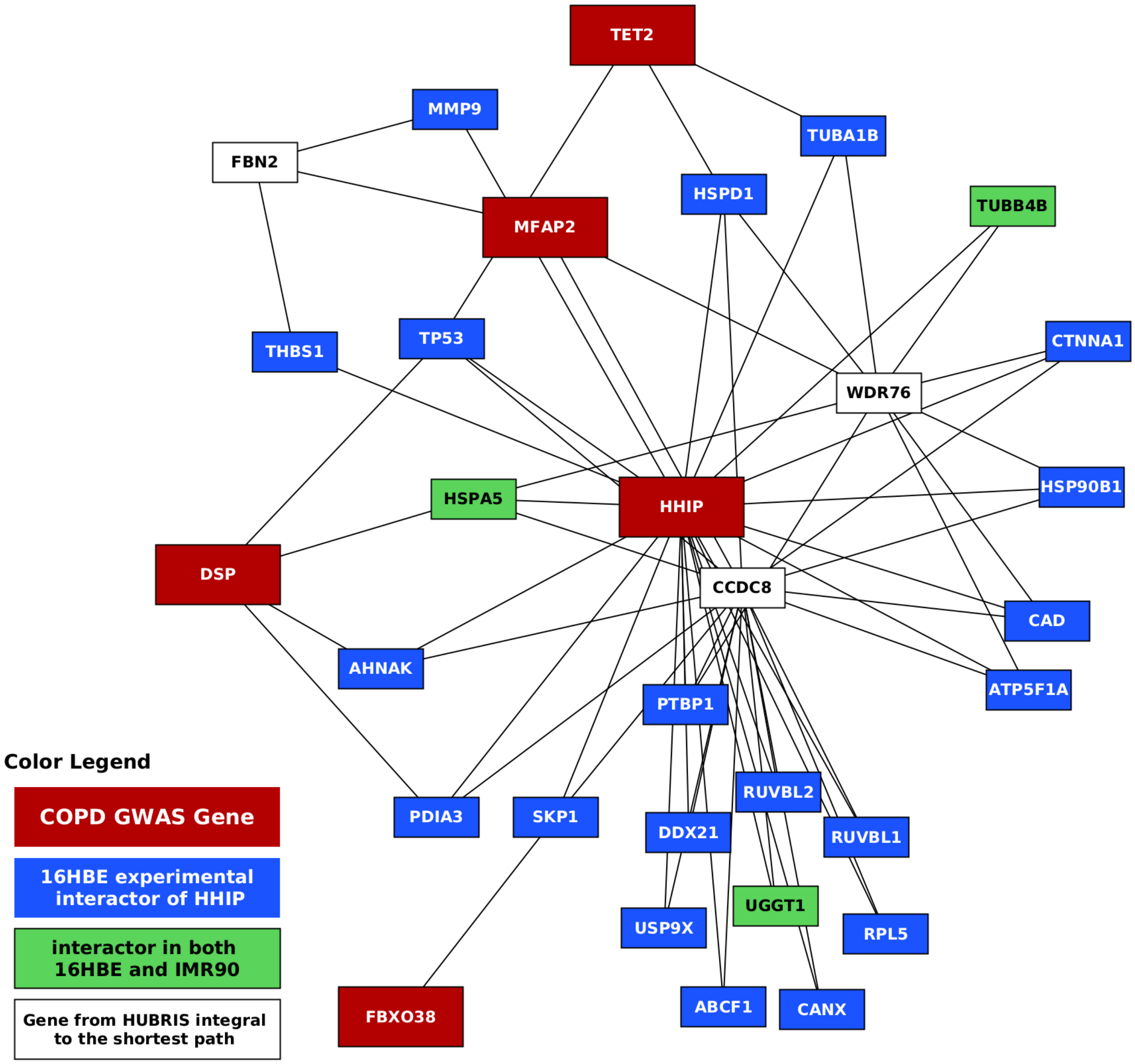


***Figure S5: New, shorter paths between HHIP and COPD GWAS genes in 16HBE.*** *The experimental links are determined in SAINTexpress (without CRAPome). The COPD GWAS genes are larger with a red color, and we also highlight nodes that were parts of shorter paths in IMR90 cells.*

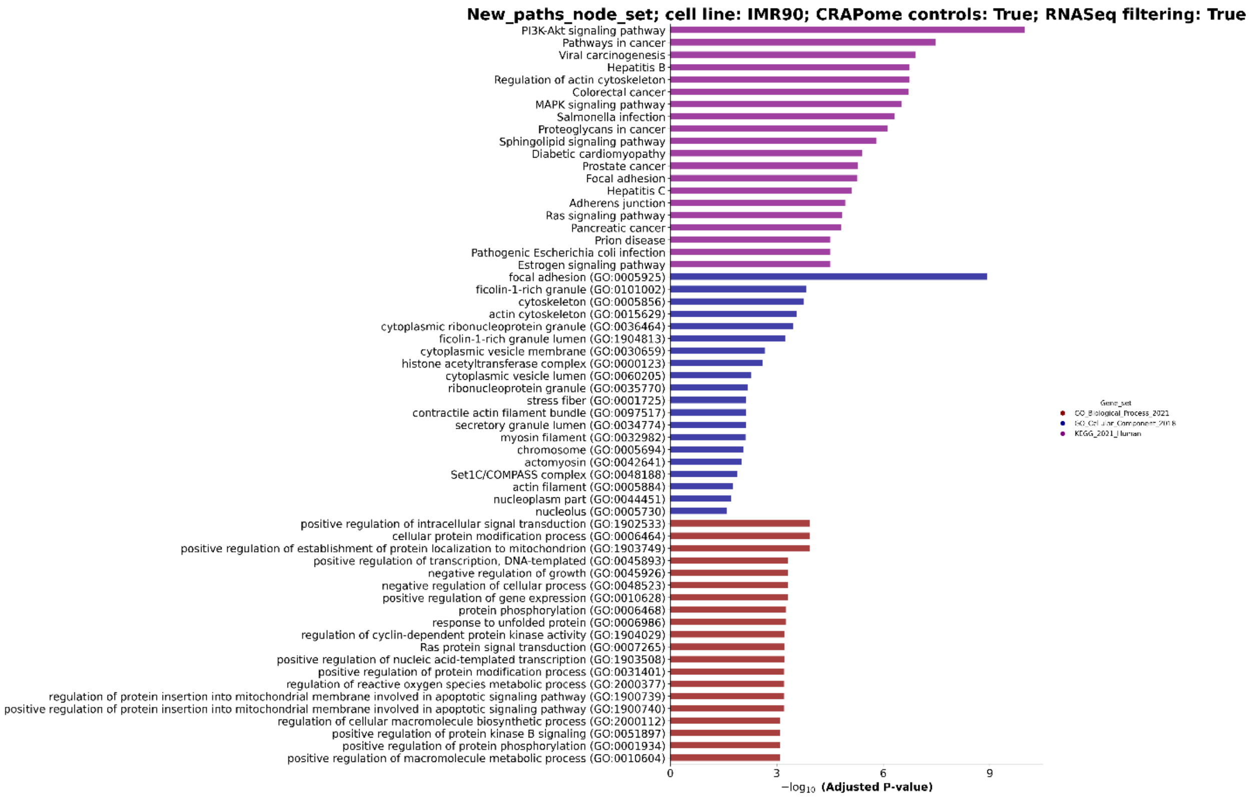


***Figure S6: Functional enrichment analysis of the nodes in new shortest paths between HHIP and the COPD GWAS genes*** *enabled by the new experimental interactions in IMR90 cells.*

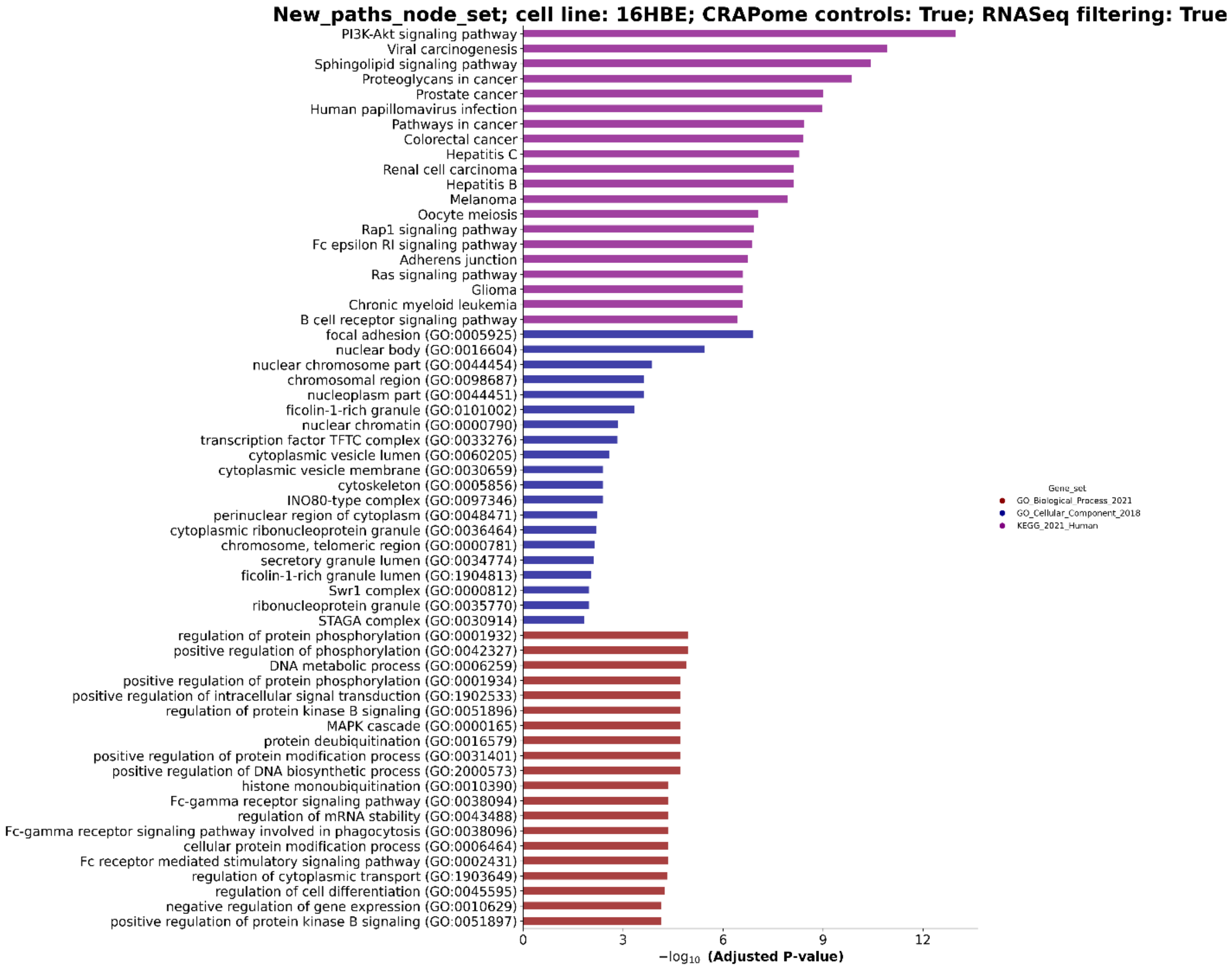


***Figure S7: Functional enrichment analysis of the nodes in new shortest paths between HHIP and the COPD GWAS genes*** *enabled by the new experimental interactions in 16HBE cells.*

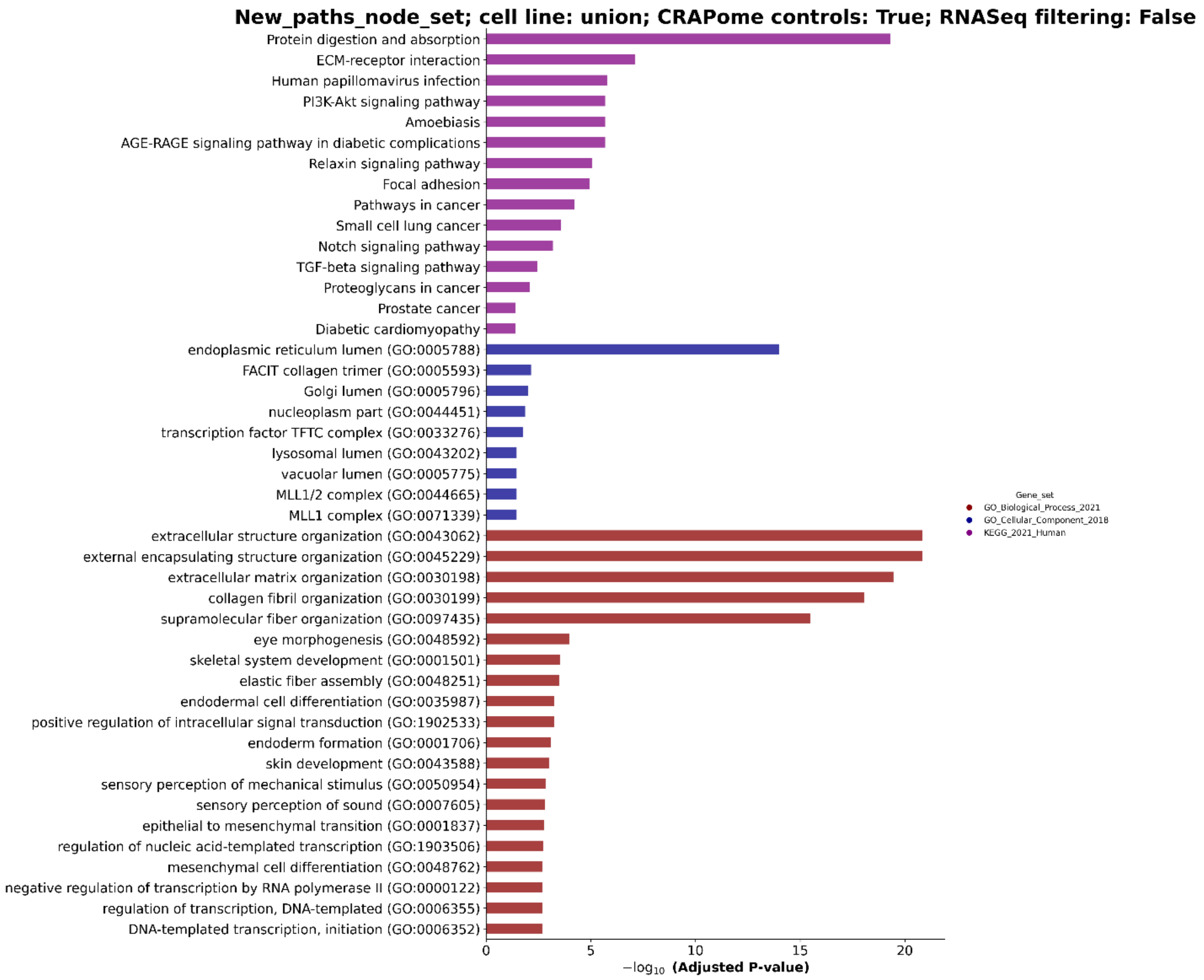


***Figure S8: Functional enrichment analysis of the nodes in new shortest paths between HHIP and the COPD GWAS genes** enabled by the new experimental interactions in both cell lines with no RNA-Seq based filtering.*

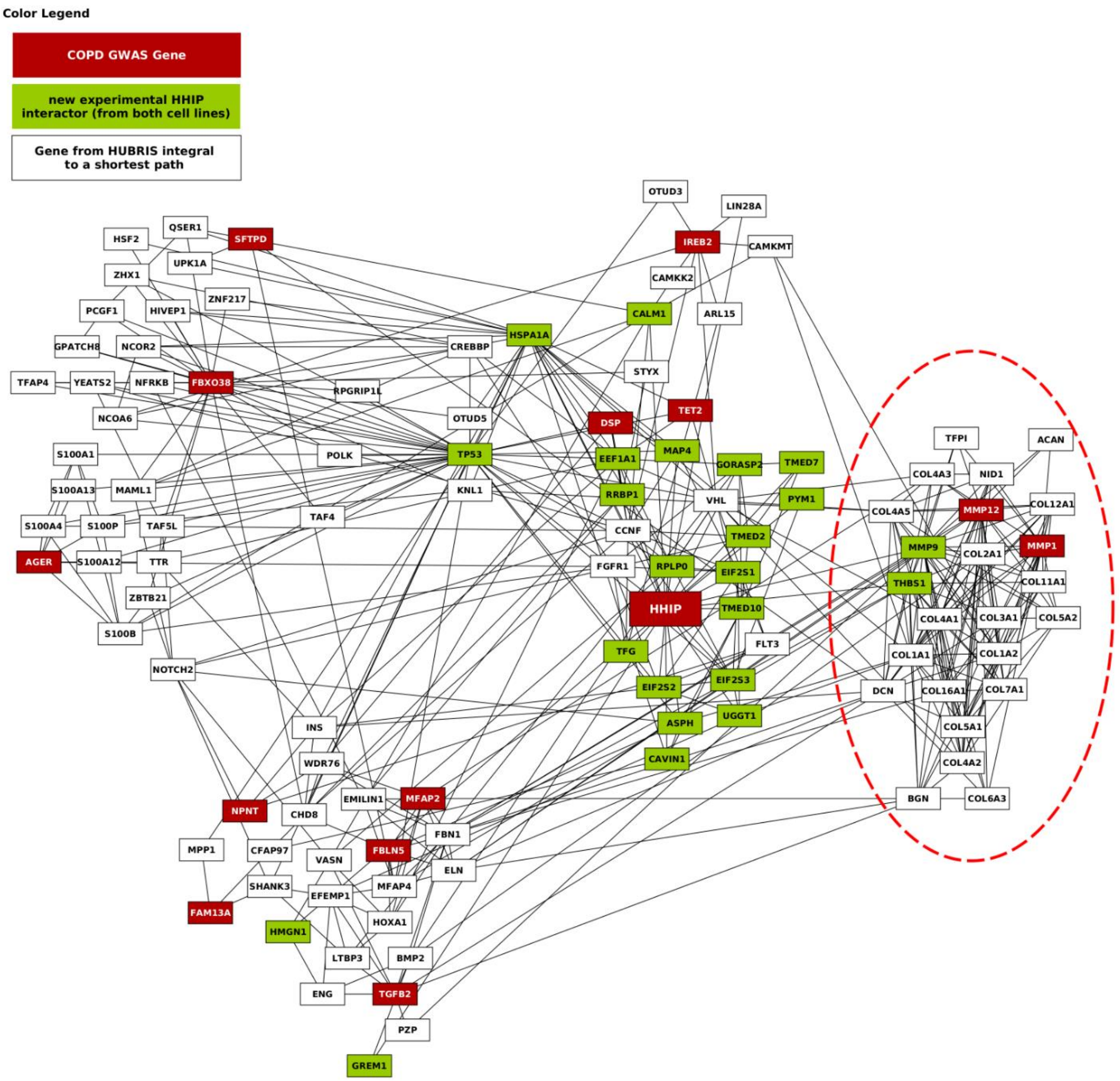


***Figure S9: Sub-network of the nodes in new shortest paths between HHIP and the COPD GWAS genes enabled by the new experimental interactions in both cell lines with no RNA-Seq based filtering.*** *The red circle highlights the cluster of collagen regulators and MMPs, which leads to the enrichment for extracellular matrix and tissue organization interactions.*

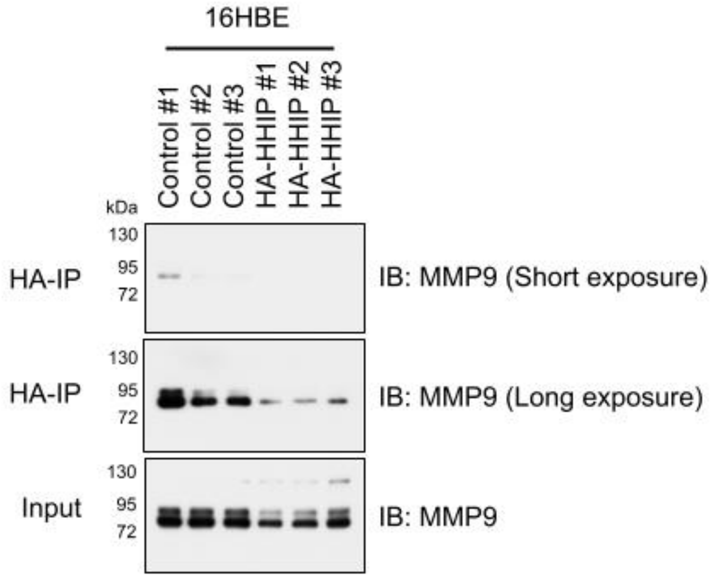


***Figure S10: Validation of the interaction between immunoprecipitated HA-HHIP and endogenous MMP9.*** *IB analysis of triplicate eluted HA-IPs and inputs derived from 16HBE cells stably expressing HA-HHIP.*

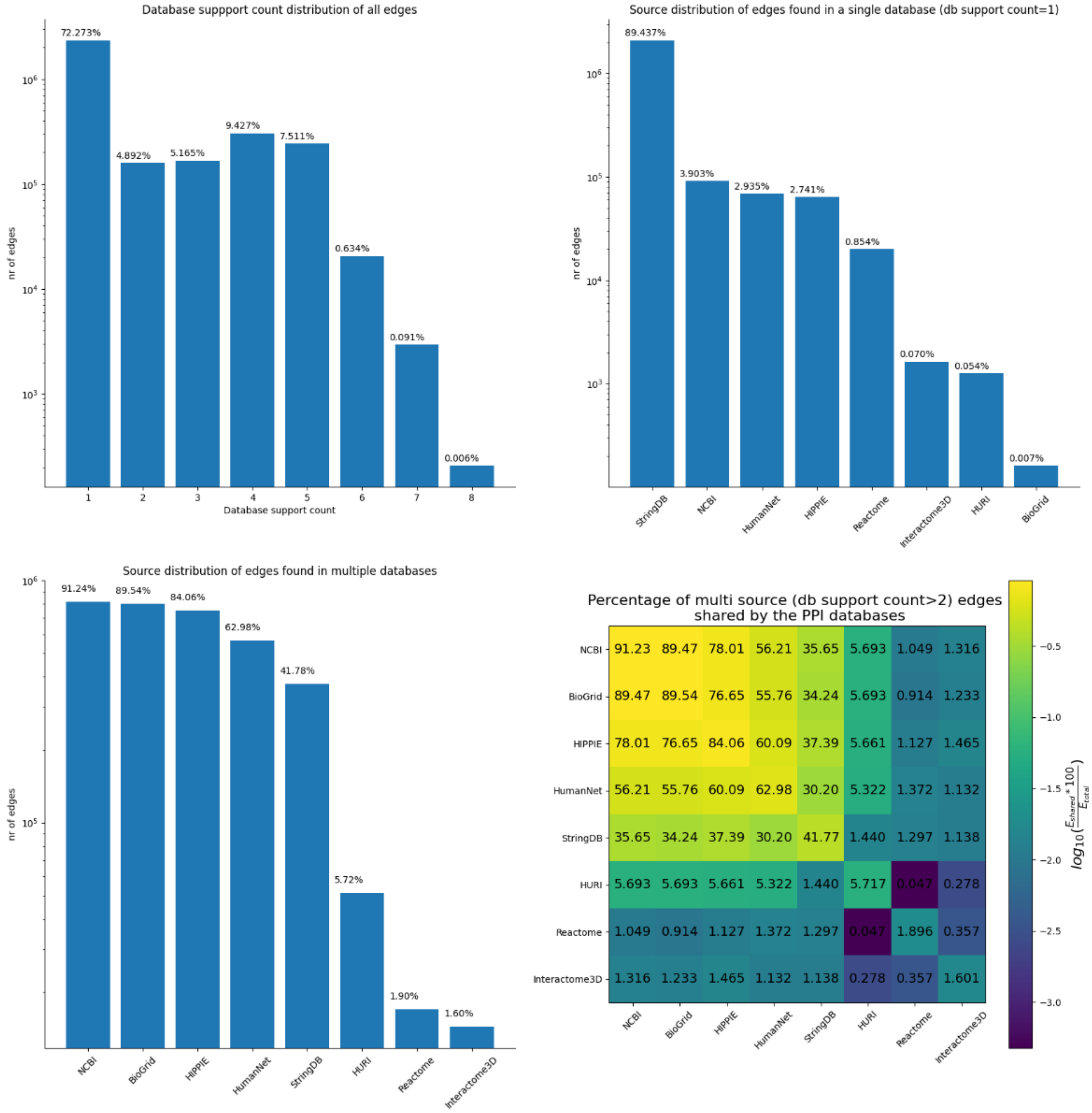


***Figure S11: Metrics on the database support of the network edges in the HUBRIS network*. *Top left*:** *Database support count for each edge in the raw HUBRIS network (simple merge without filtering). Most edges (>72%) are found in a single database.* ***Top right****: Source distribution on the subset of edges, which can be found in a single database. Most of these edges are from StringDB (>89%).* ***Bottom left****: Source distribution on the subset of edges that can be found in at least two databases. The sources are much more evenly distributed.* ***Bottom right****: percentage of edges supported by different pairs of databases, which also shows degrees of overlap between the different databases.*

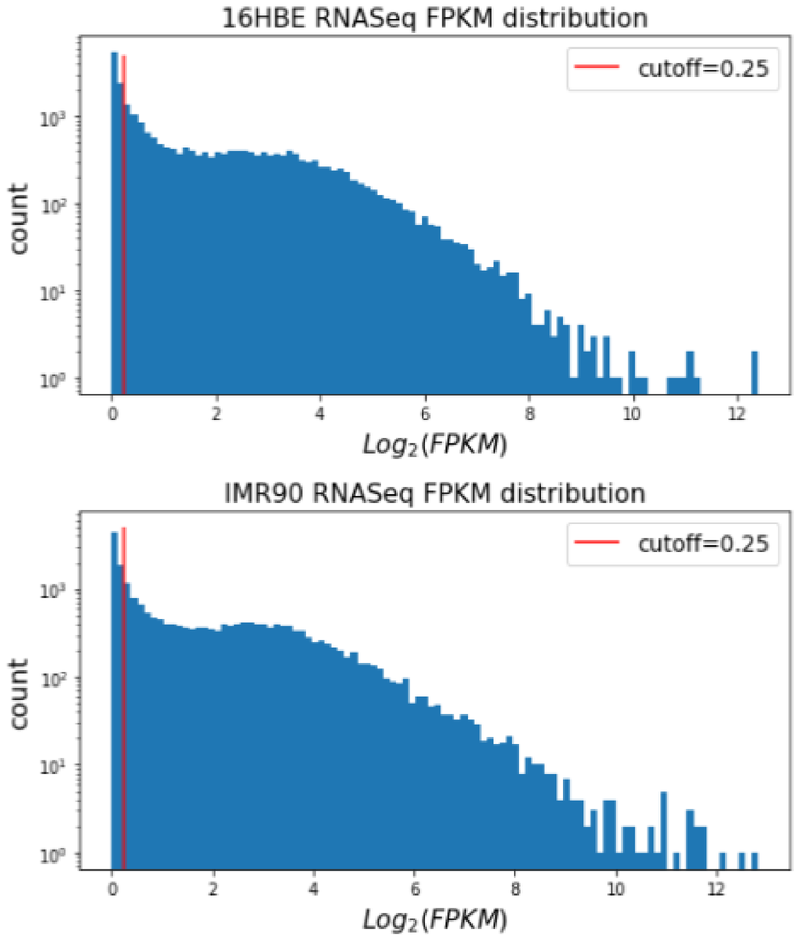


***Figure S12 Distribution of the Fragments Per Kilobase of transcript per Million mapped reads (FPKM) of our RNA-Seq data on two cell lines****: 16HBE (top) and IMR90 (bottom). The red vertical line highlights the cutoff value above which we consider genes to be expressed.*

| Gene | Assay ID | Exon Location |
|---|---|---|
| IL6 | Hs.PT.58.40336675 | 4-5 |
| IL8 | Hs.PT.58.39926886.g | 1-1 |
| IL33 | Hs.PT.58.21416460 | 2-7 |
| SIRT1 | Hs.PT.58.40382601 | 7-8 |
| SIRT3 | Hs.PT.58.22633153.g | 2-3 |

**Supplementary Table S5:** All of the probes were ordered from IDT (Integrated DNA Technologies) From PrimeTime™ Predesigned qPCR Assays.

### Legend for Supplementary Tables 1-4 (provided as separate files)

**Supplementary Table S1**: All significant protein interactors of HHIP identified by AP/MS experiments in lung cell lines. The rows of the table correspond to the different genes that are significant in at least one of the different experiments/validation pipelines. The columns represent the combination of cell line (IMR90, 16HBE), experiment (06, 11) and validation pipeline (SAINTexpress, SAINTexpress with CRAPome controls). If the value of the cell is 1 then there is a significant link between HHIP and the gene in the bait_gene_symbol column in the respective cell line/experiment/validation.

**Supplementary Table S2**: Significant pathways (FDR<0.05) of the newly identified HHIP interactor sets. The pathway column contains the pathways for which significant functional enrichment is found in at least one set. The rest of the columns represent HHIP interactor sets identified based on the combination of 3 different variables: cell line (16HBE/IMR90/union of both cell lines), the use of CRAPome controls in the SAINTexpress analysis (True/False) and the filtering of gene products based on cell type-specific RNA-Seq data (True/False). The cells contain the p-value of the enrichment for the given pathway, while "ns" means not significant. For how the different sets are derived see Methods and Supplementary Table S1.

**Supplementary Table S3**: Significant pathways (FDR<0.05) of the nodes creating the new shortest paths between HHIP and the other COPD GWAS gene products. The set of new shortest path-nodes depends on the set of new HHIP interactors we add to the HUBRIS network. The pathway column contains the pathways for which significant functional enrichment is found in at least one set. The rest of the columns represent HHIP interactor sets identified based on the combination of 3 different variables: cell line (16HBE/IMR90/union of both cell lines), the use of CRAPome controls in the SAINTexpress analysis (True/False) and the filtering of gene products based on cell type-specific RNA-Seq data (True/False). The cells contain the p-value of the enrichment for the given pathway, while "ns" means not significant. For how the different interactor sets are derived and how the new path node sets are established see Methods and Supplementary Table S1.

**Supplementary Table S4**: Significant pathways (FDR<0.05) of the nodes of the induced graph between HHIP and the other COPD GWAS gene products. The node set of the induced graph depends on the set of new HHIP interactors we add to the HUBRIS network. The pathway column contains the pathways for which significant functional enrichment is found in at least one set. The rest of the columns represent HHIP interactor sets identified based on the combination of 3 different variables: cell line (16HBE/IMR90/union of both cell lines), the use of CRAPome controls in the SAINTexpress analysis (True/False) and the filtering of gene products based on cell type-specific RNA-Seq data (True/False). The cells contain the p-value of the enrichment for the given pathway, while "ns" means not significant. For how the different interactor sets are derived and how the induced graph is defined see Methods and Supplementary Table S1.
